# Variable patterns of phenotypic evolution among canonical ‘living fossil’ lineages

**DOI:** 10.1101/2024.11.14.623638

**Authors:** Rafael A. Rivero-Vega, Jacob S. Berv, John T. Clarke, Matt Friedman

## Abstract

Coelacanths, lungfishes, and holosteans represent three emblematic living fossil lineages, thought to be united by similar patterns of phenotypic change through time. While past studies suggest that diverse evolutionary patterns occur within these groups, it is unclear whether these reflect biological differences or arise from contrasting analytical approaches. Here, we examine these lineages under a common framework to assess variation in the evolution of discrete characters, and morphometric shape data, to test whether living fossils show comparable patterns of phenotypic evolution. Our results suggest different evolutionary modes occur, both among and within lineages, as a function of data type. For lungfishes, rates in discrete characters are highest in the Devonian and monotonically decline over time. Coelacanth rates show multiple early peaks followed by a decline toward the recent. Holostean rates show modest peaks but are broadly comparable over time. Patterns of body shape evolution also differ among clades, with strong support for declining rates over time for coelacanths but mixed evidence for similar dynamics in the other groups. Our results imply idiosyncratic processes of evolutionary change among traditional examples of living fossils and indicate a need to explicitly quantify patterns of change rather than apply informal, often qualitative, macroevolutionary classifications.

## 1. Introduction

“Living fossil”—a term coined by Darwin [1]—is one of the most evocative ideas in evolutionary biology. Like many concepts first articulated in *On the Origin of Species*, the initial account of living fossils combined careful natural history observations with hypotheses of their origins. Darwin provided three examples of living fossils: ganoid fishes (bichirs, sturgeons, paddlefishes, gars, and the bowfin), *Ornithorynchus* (platypus), and *Lepidosiren* (South American lungfish) ([1]: p. 107). These groups were, Darwin argued, united by a set of common features. First, each contained the last survivors of lineages that the fossil record indicates were once much more diverse. Second, all three represented structural intermediates between more species-rich groups (chondrichthyans and teleosts in the case of ‘ganoids’, reptiles and mammals in the case of *Ornithorhynchus*, and fishes and tetrapods in the case of *Lepidosiren*). Third, all are restricted to specific, often spatially limited habitats. Darwin saw in this final observation a potential explanation for the first two features of living fossils: endurance of such lineages might reflect reduced competition encountered in geographically or environmentally restricted settings.

Since Darwin’s proposal, the concept of living fossils has been widely applied across the tree of life [2]. As the set of lineages interpreted as living fossils has expanded, the concept’s meaning has grown so diffuse that its utility is hotly debated [3–9]. Investigations into the dynamics of phenotypic evolution reflect a shift from Darwin’s original formulation of living fossils to more recent conceptualisations [10,11]. Although Darwin only indirectly addressed processes of morphological change in these groups, aspects of evolutionary tempo and mode responsible for patterns of phenotypic variation are central to many of today’s applications of the living fossil concept (e.g., see criteria in [7]). Generally, the phenotypic variation in living fossil groups is interpreted as either uniformly low [12], subject to strong evolutionary constraint [13], or declining over time [14,15], with the latter being more prevalent in the recent literature (e.g., *Sphenodon* [15]). With some exceptions [16], quantitative investigations of phenotypic evolution in putative living fossil lineages are restricted to isolated examples. While some aspects of tempo and mode may be shared, assessing these lineages independently prevents fair comparisons among them [14,17–21].

To assess the degree of macroevolutionary convergence in aspects of phenotypic evolution among the most prominent examples of living fossils, we combined existing and newly collected morphological data and analysed each taxon in a consistent framework. Our goal is not to test *a priori* expectations of specific macroevolutionary patterns. Instead, we look to test whether our canonical living fossil groups share comparable patterns of phenotypic change over time. Such similarity is implied by much of the recent literature on living fossils [14,15]; however, it is not reflected in Darwin’s three defining characteristics of such groups. In essence, we are testing whether the gross patterns that unite living fossils might be generated by similar underlying histories of evolutionary change. This approach is similar to that of “phylogenetic natural history” *sensu* Uyeda et al. 2018 [22], with the aim of documenting phylogenetic evidence of evolutionary process. Importantly, we do not seek a numerical measure of potential “living fossil”-ness like some recent work [23]. Rather, our methodological approach is inspired by recent cross-clade comparative approaches used to evaluate adaptive radiations [24,25], another macroevolutionary concept that had come to be broadly—and at times imprecisely—applied since it was first articulated [24,26–30]. We take a restricted approach, emphasising two of the lineages first identified by Darwin: lungfishes and holosteans. Darwin’s third example, monotremes, are too scarce in the fossil record [31] to offer a helpful comparison, so we examine coelacanths instead. Though obviously not considered by Darwin because eight decades separate the publication of *Origin* from the naming of the living *Latimeria*, coelacanths conform to the original definition of living fossils. Features common to these three groups make them ideal for the comparative study of phenotypic evolution: a long history of systematic and paleontological examination, established phylogenetic frameworks, and articulated fossils that provide perspectives on changes in gross morphology over time. Prior studies of evolutionary rates in holosteans [32], lungfishes [14,17,33], and coelacanths [18–20,33] focus on different aspects of phenotype using contrasting analytical tools that limit comparisons. Here, we examine the evolution of discrete (i.e., cladistic characters) and continuous (i.e., body shape) traits in these three groups in a common framework to address whether these examples of living fossils show similar patterns of phenotypic change over time.

## 2. Materials and Methods

### (a) Discrete Character Matrices

We used published morphological character matrices to simultaneously infer time-scaled phylogenetic hypotheses and rates of discrete-trait evolution in coelacanths, lungfishes, and holosteans (see Phylogenetic Analyses). Our analyses of shape evolution within a model-fitting framework then used these phylogenetic hypotheses. The source character matrices are described below. For all matrices, we collapsed ambiguous or polymorphic character states to question marks (?) prior to running new phylogenetic analyses.

For coelacanths, we used the matrix from Toriño et al. [34], a descendant of the matrix presented by Forey [18] and one which has been used, with modest modifications, in a series of additional studies [35–37]. The matrix contains 50 taxa (48 coelacanths) scored for 110 unordered characters, 87 of which are cranial and 23 of which are postcranial (inclusive of scales).

For lungfishes, most character matrices focus on either Devonian [38,39] or post-Devonian [40,41] taxa, with few offering dense taxonomic sampling over the entire evolutionary history of the group [14,21,42,43]. From the available matrices, we selected Lloyd et al. [14], which represents a ‘supermatrix’ combining datasets from several studies [38–40,44]. This matrix formed the basis of a relatively recent assessment of evolutionary rates in lungfishes [14], permitting direct comparison with our results obtained from a different analytical approach. This matrix contains 86 taxa (85 lungfishes) scored for 91 characters, 82 of which are cranial and 9 of which are postcranial (inclusive of scales).

For holosteans, we used the dataset from López-Arbarello and Sferco [45] that includes a broad sample of holosteans and other crown neopterygians. It contains 70 taxa (39 holosteans) scored for 339 characters, 192 of which are cranial and 145 of which are postcranial (inclusive of scales and dermal elements).

### (b) Phylogenetic Analyses

We performed bayesian phylogenetic analyses for the lungfish, coelacanth, and holostean morphological datasets in BEAST v2.6.5 [46,47]. We applied the Fossilized Birth-Death (FBD) approach, which uses extinct taxa as informative tips and simultaneously estimates relationships, branch durations, and rates of morphological change through time [48,49]. Empirical studies and simulations show that use of the FBD prior generally results in estimated divergence times that are more closely in agreement with the fossil record over other tree inference models [50–53]. We used a Lewis MK model [54] with 4-category gamma distributions and a relaxed lognormal clock for morphological traits. Relative to the default settings in BEAST, we changed two priors specifically for coelacanths and one additional prior across all groups. In coelacanths, the two priors were the *diversificationRateFBD* (Log Normal; upper = 10) and the *ucldMean* (Log Normal distribution; initial = 0.1; M = 1.0; S = 1.25). Across all three groups, the *originFBD* prior was the only additional change. For both sarcopterygian lineages, we set the initial search to 425 Ma (Uniform distribution, upper bound of 440 Ma), which we selected based on the age range estimates of the most recent common ancestor (MRCA) between sarcopterygians and actinopterygians (*Guiyu* from the late Silurian [55]). For holosteans, we set the initial search to 260 Ma (Uniform distribution, upper bound of 423 Ma), which we selected based on the MRCA for the group. All remaining FBD priors apart from those specified individually per clade above were left in their default settings—Uniform for the *diversificationRateFBD* (initial = 0.01; upper = infinity), *samplingProportionFBD* (initial = 0.5; upper = 1.0), *turnoverFBD* (initial = 0.5; upper = 1.0), and *ucldMean* (initial = 1.0; upper = infinity); Exponential for the *gammaShape* (initial = 1.0; mean = 1.0); and Gamma for the *ucldStdev* (initial = 0.1; alpha = 0.5396; beta = 0.3819). We did this to reduce biases associated with the *ad hoc* adjustment of informative priors by limiting their use to “realistic” mechanistic models at the root search [56].

We applied a small set of constraints to our analyses. Topological constraints included: (1) rooting using a designated outgroup (actinopterygians for coelacanths, *Psarolepis* for lungfishes, and *Pteronisculus* for holosteans), (2) monophyly of the apomorphic Paleozoic coelacanths *Allenypterus* and *Holopterygius* [35], (3) monophyly of a well-supported subset of total-group Lepidosireniformes for lungfishes (*Gosfordia truncata*, *Lepidosiren paradoxa*, *Protopterus annectens*, and *Ptychoceratodus serratus*) [41]. Temporal constraints included: (1) a fossil calibration minimum at 100 Ma for the node representing the MRCA of *Lepidosiren paradoxa* and *Protopterus annectens* based on the breakup of Gondwana [57–59] and (2) a calibration setting a minimum age of divergence between *Lepiososteus* and *Atractosteus* at 93 Ma, based on the first occurrence of fossil *Atractosteus* in the early Late Cretaceous [60]. We ran the phylogenetic analyses for 500 million generations, with trees sampled every 100,000 generations. We used Tracer v1.7 [61] to assess convergence visually via the trace plot and effective sample size (ESS) calculations. We then used the TreeAnnotator utility packaged with BEAST to ignore the first 10% of the trees sampled as burn-in fraction and use the post burn-in trees to generate MCC consensus trees (leaving the full posterior sample files unedited). Because our downstream analyses considered only those tips assigned to each total group of interest (e.g., coelacanths, lungfishes, and holosteans), we pruned other clades from both the MCC tree and the trees sampled from the posterior.

We compiled age constraints for fossil taxa at the finest level possible (typically geologic stage [62–64]), drawing from a combination of original descriptions, revisions, and other resources (e.g., The Paleobiology Database, https://paleobiodb.org/). In BEAUTi v. 2.6.5 [47], the age prior for each extinct taxon was set as a uniform distribution. The upper and lower boundaries are defined by the age estimates for the most restrictive stratigraphic bin identified for each fossil taxon. Likewise, if there was more than one fossil from different deposits or localities representing the taxon, we used the most restrictive bin that accounted for its overall stratigraphic extent. We then linked age priors for taxa deriving from the same geological horizon (typically a well-known fossil site or formation) using the R package *palaeo* [65] to generate an XML code block that would be appended manually to the BEAUTi input file used to initialise BEAST. This enforced matching ages for fossils from the same deposits when running phylogenetic analyses and represents a novel incorporation of stratigraphic uncertainty in a comparative framework.

### (c) Evolution in Discrete Characters

Any outgroup taxa used to root the trees during the BEAST runs (including all Teleostei for the holostean tree) were excluded from summaries of evolutionary rate in discrete characters. All subsequent steps in the rate analysis pipeline used the packages *phytools* [66]*, ape* [67], and *coda* [68], along with an adjusted script from Close et al. [69] calling an additional function from *OutbreakTools* [70] in R v3.6.1 [71]. We began by reading the entire posterior distribution of trees generated during the BEAST analysis into R, removing 10% of trees as burn-in, taking a random sample of 100 trees, confirming ESS values, and then estimating per-branch rates of phenotypic evolution from this subset of trees. We plotted branches from each of the sampled trees according to their lengths (horizontal axis, millions of years) and rates (vertical axis, natural log scale) to depict a “rate cloud”. We then imported the MCC tree and used it to add a generalised loess line by calculating a local average rate across lineages in million-year steps. Our overall depiction, therefore, represents the average natural log rate change across time for the entire clade ([69]: figure 4).

In addition to variation in rates of change over time, we evaluated character saturation levels within all three groups using their respective morphological matrices. Prior research has investigated saturation to estimate if convergence in character states is more likely than the generation of new character states [72–75]. This was achieved by comparing pairwise character-state dissimilarity and plotting it against patristic morphological distance (*distance_metric* = “mord”) [76]. In these analyses, if the average value (loess curve) increases linearly, it indicates persistent generation of novel traits. However, if the average value asymptotes, it signals lineages reaching character exhaustion with increased incidence of homoplasy. These analyses illustrate the relative contributions of innovation and homoplasy to overall change. Although there may be limitations in this method due to the coding of autapomorphic characters, potential mitigations would require estimation of saturation from model-based inferences of character states, which is beyond the scope of the present study.

The code and data used to run all subsequent analyses will be available on GitHub and archived on Dryad upon acceptance of the manuscript.

### (d) Morphological Landmarking Schemes

We selected landmarks based on previous publications with adjustments made on a per-clade basis to capture homologous aspects of cranial and postcranial anatomy (see clade information below). All semi-landmarks were spread out evenly beginning at the rostrum, following the dorsal body line including the fleshy part of any fins, curving at the fleshy posterior end of the caudal fin, and then proceeding ventrally while tracing the fleshy part of any additional fins, and then ending at the posteriormost contact of the lower jaw with the ventral body line. We did not use a consistent number of semi-landmark points between fixed landmarks; rather, the total semi-landmarks varied depending on the topological complexity of the region they were outlining. Non-identical constellations of fixed landmarks for the three lineages reflected important differences in the number and arrangement of median fins. The full landmarking schemes for each clade are discussed below and figured in the electronic supplementary material.

The arrangement of fixed landmarks for coelacanths matches that used by Friedman and Coates [35], with 8 landmarks capturing fin position and 6 landmarks capturing cranial features (14 total, electronic supplementary material, figure S4). 155 additional semi-landmarks were placed to capture the outline of the body along segments beginning at the anteriormost tip of the skull and tracing the head (one curve, 25 semi-landmarks); the dorsal region, including tracing around the fleshy area of each fin lobe (four curves, 35 semi-landmarks); the fleshy dorsal and ventral regions of the caudal fin (two curves, 60 semi-landmarks); and the entire ventral region, including tracing around the fleshy area of each ventral midline fin, ending posteriorly at the lower jaw (two curves, 35 semi-landmarks).

The landmarking scheme for lungfishes was built to closely match the configuration of the coelacanth scheme, resulting in 5 landmarks capturing fin position and 6 landmarks capturing variations in skull morphology (11 total, electronic supplementary material, figure S5). The reduced number of fixed cranial landmarks reflects the absence of median fin divisions in all lungfishes; these divisions remain apparent in coelacanths with the most divergent postcranial anatomy [35]. 115 additional semi-landmarks were placed to capture the outline of the body along segments beginning at the anteriormost tip of the skull and tracing the head (one curve, 15 semi-landmarks), the entire dorsal region, including tracing around the fleshy area of each fin lobe, extending to the posterior tip of caudal fin (two curves, 50 semi-landmarks), the entire ventral region, including tracing around the fleshy area of each ventral midline fin, ending at the lower jaw (two curves, 50 semi-landmarks).

The landmarking scheme for holosteans descends from that used in Clarke et al. [77] and Clarke and Friedman [32], with 8 landmarks capturing fin position and 6 landmarks capturing variation in skull morphology (14 total, electronic supplementary material, figure S6). 120 additional semi-landmarks were placed to capture the outline of the body along segments beginning at the anteriormost tip of the skull and tracing the head (one curve, 25 semi-landmarks); the dorsal region, including tracing around the fleshy area of each fin (four curves, 55 semi-landmarks); the fleshy dorsal and ventral regions of the caudal fin (two curves, 20 semi-landmarks), and the entire ventral region, including tracing around the fleshy area of each ventral midline fin, ending posteriorly at the lower jaw (one curve, 20 semi-landmarks).

### (e) Body Shape Analyses

We landmarked a combination of original specimen photographs, published fossil images, and technical illustrations for species that were present in our phylogenetic analyses. The final dataset consisted of 27 coelacanths, 16 lungfishes, and 62 holosteans. Each taxon was then landmarked according to the schemes outlined above and their body shape was traced with spline curves in R v3.6.1 [71] using the package *StereoMorph* (*digitizeImages*) [78]. The unaligned landmark and semi-landmark coordinates were imported using the package *geomorph* [79,80] (*readland.shapes*, adapting the total semilandmarks on a per-curve basis to be dense enough to capture fin details as detailed above), adjusted with a Procrustes transformation (*gpagen*), and transformed using a principal components analysis (PCA) and phylogenetically-aligned components analysis (PaCA) (both via *gm.prcomp*). Morphospace plots for all taxa can be found in electronic supplementary material figures S4–S6.

### (f) Fitting Evolutionary Models of Continuous Trait Evolution

We fit three alternative models of continuous trait evolution to body shape datasets for each clade using the R package mvMORPH [81], taking advantage of new approaches for investigating high-dimensional morphometric datasets [82]. For each clade, we fit: (1) Brownian Motion (BM), (2) Ornstein-Uhlenbeck (OU), and (3) Accelerating-Decelerating (ACDC) models. BM reflects a constant accumulation of shape disparity over time; OU models a central tendency, which is interpreted biologically as an adaptive peak or evolutionary constraint [83,84]; and ACDC reflects accelerating or decelerating exponential rates of change over the history of a clade [81,85]. Early Burst (EB) is a special case within the ACDC framework where rates exclusively decelerate [24,81,85]. EB was originally formulated as a model of adaptive radiation [24], but it also matches patterns sometimes inferred for living fossils [14,17]. For the purposes of our analyses, we refer to the exponential models inferring declining rates as EB.

We fit the three alternative models using the *mvgls* function in mvMORPH. We opted for the penalised likelihood approach in *mvgls* because it is designed to operate directly on highly dimensional morphometric datasets and can efficiently optimise multivariate models with hundreds of variables [82]. Additionally, other likelihood models can lead to, for example, exaggerated estimates of the exponential parameter *b* in likelihood (e.g., *mvEB*) versus penalised likelihood (e.g., *mvgls*) model fitting analyses [82]. The *mvgls* flags we used across all analyses were *scale.height* = F, *method* = “PL-LOOCV”, *penalty* = “RidgeArch”, *target* = “unitVariance”, and *error* = T. The error option estimates an extra nuisance parameter intended to account for variation not explained by the model (e.g., measurement error) and is recommended for empirical datasets [82,86].

To run the analyses, we used the MCC tree as an initial phylogenetic framework and all PC axes— those explaining 100% of the variation—for each taxon (24, 15, and 60 total axes for coelacanths, lungfishes, and holosteans, respectively). To account for variation in model fit as a function of uncertainty in tree topology and evolutionary timescale, we repeated the procedure using a sample of 100 trees from the posterior distribution of trees generated by the BEAST analyses. We calculated GIC scores [82,87,88] and weights for each model fit across all sampled trees, and visualised the resulting distributions (figures 1*c*, 2*c*, 3*c*; GIC scores in the electronic supplementary material).

**Figure 1.**
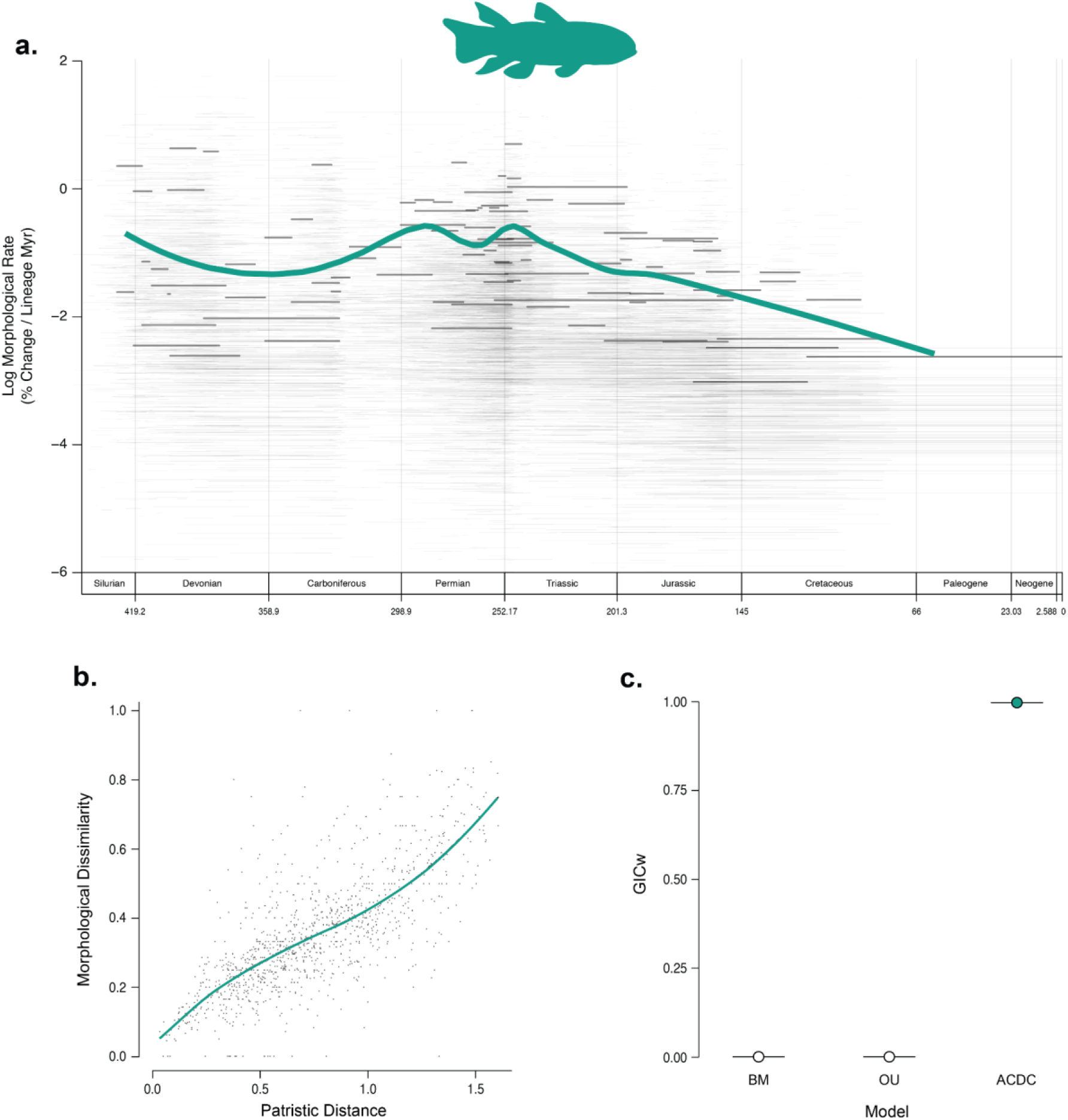
Patterns of phenotypic evolution in coelacanths. (a) rate-through-time plot for discrete characters, derived from the MCC tree and a random sampling of 100 trees from the posterior generated using morphological data in BEAST v2.6.5 with the Fossilized Birth-Death model. The thicker, solid green line is the average rate across the MCC tree; the thinner, solid black lines are the MCC per-branch rates; and the semi-transparent grey lines forming a “cloud” behind the plot are the per-branch rates from each tree sampled from the posterior. Average rate-through-time plots for all groups can be found in Figure S9. (b) character saturation, generated using the discrete character matrix to calculate pairwise character-state dissimilarity and plot it against patristic morphological distance. (c) support for competing models of body-shape evolution (GICw) derived from the MCC tree (larger circles), a sample of 100 posterior trees (violin plot distribution with smaller circles), and the PC axes that summarised 100% of the variability for each clade. The large, coloured circle indicates the highest model support recovered for the MCC tree. The black box plots within the violin plots have smaller, grey circles denoting the median model support across the 100 posterior trees tested. The intervals represented by the boxes and whiskers are the quartiles calculated by the *vioplot* package. Models and *mvMORPH* methods are discussed in the text. Silhouette of *Miguashaia* by Steven Coombs (Public Domain) via PhyloPic.

We also performed parametric bootstrapping—using only the parameters from the best fit model per clade—to explore our results more thoroughly. The empirical parameters for the *mvSIM* function were thus derived from *mvgls* model fitting using the unit-scaled MCC tree and the PC axes explaining 100% of the variance. Due to optimization issues, holosteans required the *mvgls* parameter under *model* = “EB” to be estimated using *REML* = FALSE throughout. Coelacanths and lungfishes showed no optimization issues and used the default *REML* = TRUE. We then performed bootstrapping on 100 *mvSIM* replicates, with the upper and lower bounds of parameter search space iteratively increased if model fitting reached the default parameter limits. We plotted the distribution of the parameter for each best fit model estimated from these bootstrapped datasets per group (electronic supplementary material, figure S8) and calculated the 95% confidence interval.

## 3. Results

### (a) Phylogenetic Relationships and Evolutionary Timescales

Our Bayesian analyses using a Fossilised Birth-Death prior produced phylogenetic topologies that correspond closely to those reported from prior analyses of the same matrices. Our estimated ages of last common ancestor of all sampled lineages for each of our groups are: Silurian for coelacanths (428 Ma; 95% HPD: 415–446 Ma; electronic supplementary material, figure S1) and lungfishes (432 Ma; 95% HPD: 425–435 Ma; electronic supplementary material, figure S2), and early Permian for holosteans (288 Ma; 95% HPD: 272–307 Ma; electronic supplementary material, figure S3). Most nodes within the holostean and coelacanth phylogenies have relatively high clade credibility values (>75%). Posterior support for most nodes in the lungfish tree is considerably lower, reflecting a longstanding pattern of heterogeneous topologies recovered in phylogenetic studies of the group (e.g., [42]). Further discussion and complete maximum clade credibility (MCC) trees can be found in the electronic supplementary material.

### (b) Discrete Character Evolution

For coelacanths, rates of discrete character evolution show an overall decline from the origin of the clade to the present (figure 1*a*). Modest peaks in evolutionary rates occur during the Early Devonian, Permian, and Early Triassic, with rates dropping substantially by the end of the Triassic. Rates plateau during the Jurassic before dropping precipitously by the end of the Cretaceous. A nearly linear relationship between morphological disparity and patristic distances indicates little character saturation (figure 1*b*).

For lungfishes, rates of discrete character evolution show an overall decline from the origin of the clade to the present (figure 2*a*). The highest rates occur in the Silurian and Early Devonian, but these decline into the Late Devonian. At this point, they stabilise for the duration of the Paleozoic. Rates decline monotonically throughout the Mesozoic and Cenozoic. An apparent asymptote in values of morphological dissimilarity with increasing patristic distance indicates some saturation of character states (figure 2*b*).

**Figure 2.**
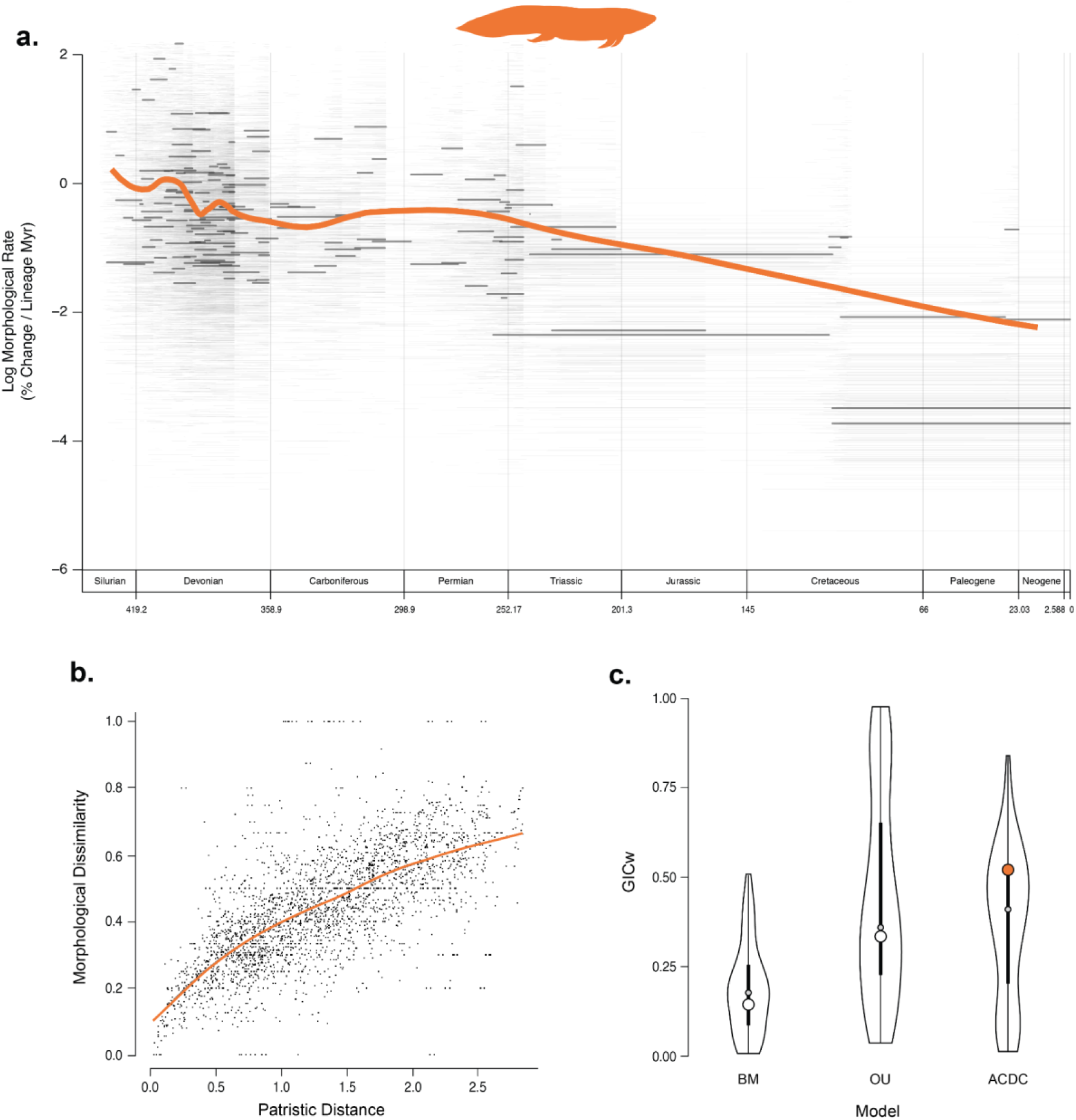
Patterns of phenotypic evolution in lungfishes. (a) rate-through-time plot for discrete characters, derived from the MCC tree and a random sampling of 100 trees from the posterior generated using morphological data in BEAST v2.6.5 with the Fossilized Birth-Death model. The thicker, solid orange line is the average rate across the MCC tree; the thinner, solid black lines are the MCC per-branch rates; and the semi-transparent grey lines forming a “cloud” behind the plot are the per-branch rates from each tree sampled from the posterior. Average rate-through-time plots for all groups can be found in Figure S9. (b) character saturation, generated using the discrete character matrix to calculate pairwise character-state dissimilarity and plot it against patristic morphological distance. (c) support for competing models of body-shape evolution (GICw) derived from the MCC tree (larger circles), a sample of 100 posterior trees (violin plot distribution with smaller circles), and the PC axes that summarised 100% of the variability for each clade. The large, coloured circle indicates the highest model support recovered for the MCC tree. The black box plots within the violin plots have smaller, grey circles denoting the median model support across the 100 posterior trees tested. The intervals represented by the boxes and whiskers are the quartiles calculated by the *vioplot* package. Models and *mvMORPH* methods are discussed in the text. Silhouette of *Ceratodus* by T. Michael Keesey (Public Domain) via PhyloPic.

For holosteans, rates of discrete character evolution show irregular but modest fluctuations about a relatively constant rate over time (figure 3*a*). Distinct peaks in rates of morphological evolution were recovered in the Triassic and Early Cretaceous, each followed by a nearly equal and opposite decrease in rates. From the Late Cretaceous onward, rates remain approximately constant. Dissimilarity increases sharply as the patristic distance increases at relatively low patristic distances. Slower accumulation at higher patristic distances indicates some character saturation (figure 3*b*). Discrete patterns are summarised for all three lineages in electronic supplementary material, table S1.

**Figure 3.**
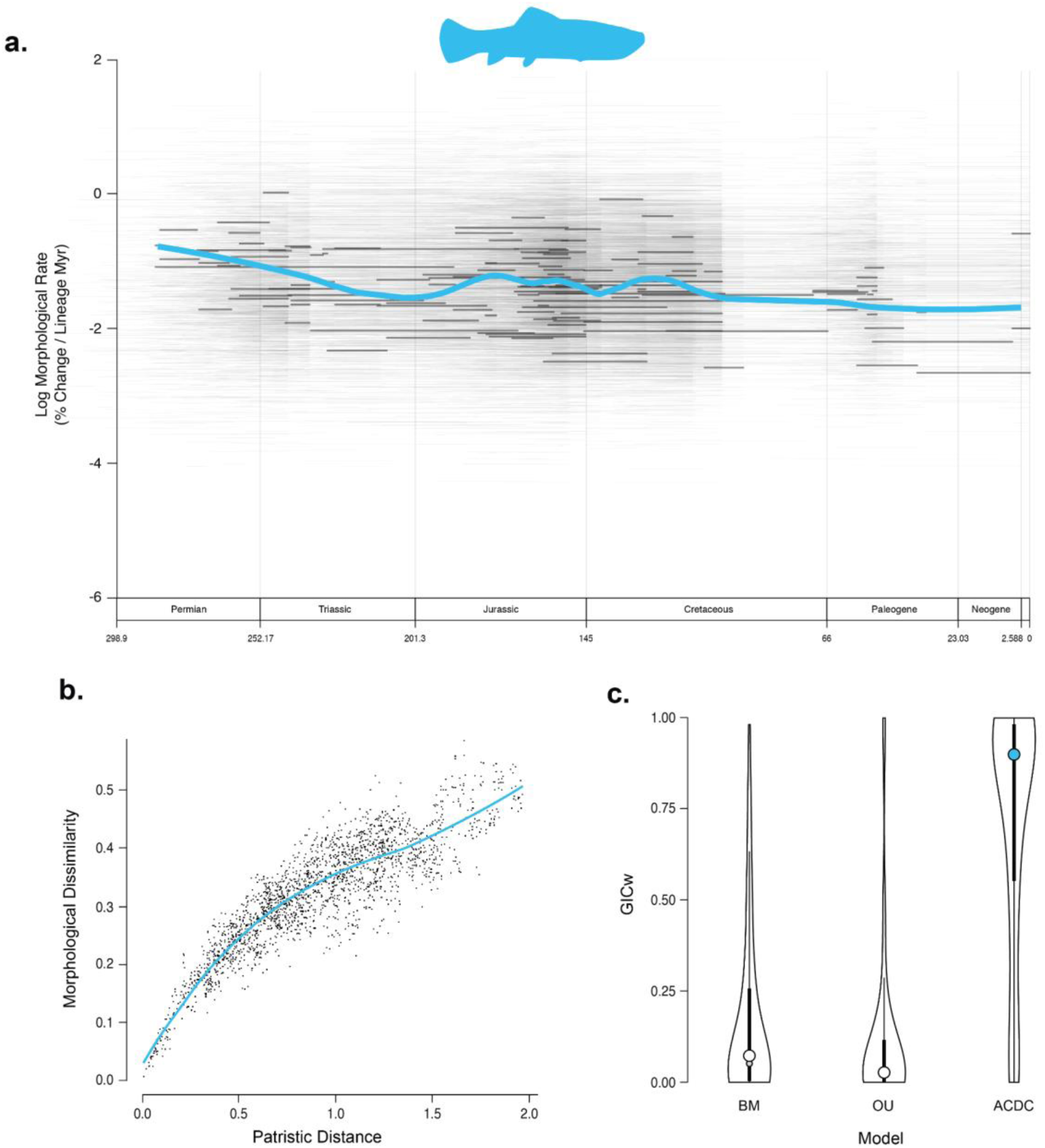
Patterns of phenotypic evolution in holosteans. (a) rate-through-time plot for discrete characters, derived from the MCC tree and a random sampling of 100 trees from the posterior generated using morphological data in BEAST v2.6.5 with the Fossilized Birth-Death model. The thicker, solid blue line is the average rate across the MCC tree; the thinner, solid black lines are the MCC per-branch rates; and the semi-transparent grey lines forming a “cloud” behind the plot are the per-branch rates from each tree sampled from the posterior. Average rate-through-time plots for all groups can be found in Figure S9. (b) character saturation, generated using the discrete character matrix to calculate pairwise character-state dissimilarity and plot it against patristic morphological distance. (c) support for competing models of body-shape evolution (GICw) derived from the MCC tree (larger circles), a sample of 100 posterior trees (violin plot distribution with smaller circles), and the PC axes that summarised 100% of the variability for each clade. The large, coloured circle indicates the highest model support recovered for the MCC tree. The black box plots within the violin plots have smaller, grey circles denoting the median model support across the 100 posterior trees tested. The intervals represented by the boxes and whiskers are the quartiles calculated by the *vioplot* package. Models and *mvMORPH* methods are discussed in the text. Silhouette of *Calamopleurus* by Aline M. Ghilardi (<u>CC BY-NC 3.0</u>) via PhyloPic.

### (c) Body Shape Evolution

Landmarking schemes and shape spaces for the three lineages are summarised in electronic supplementary material, figures S4–S6. For coelacanths, 24 principal component (PC) axes summarised 100% of body shape variance, with over 70% captured by the first two axes. PC1 reflects variation in the anterior extension of the epichordal/hypochordal lobes of the caudal fin (48% of total variance) and PC2 summarises differences in body depth (23% of total variance) (electronic supplementary material, figure S4). PC3 and PC4 capture head shape, pelvic fin placement, and posterior body shape (14% of total variance, cumulatively). All further axes summarise minor variations relating to the median fins. The best-fit model on the MCC tree was the Accelerating-Decelerating (ACDC) model with declining rates of change, corresponding to the Early Burst (EB) variant (Generalised Information Criterion [82,87,88] weight [GICw] = 1) (figure 1*c*; electronic supplementary material, figure S7, table S2). Estimated *b*—representing the exponential change in evolutionary rate—for empirical data on the unit-scaled MCC tree was strongly negative (*b* ∼ −10.7; 95% CI from parametric bootstrapping: −20.3, −8.99; electronic supplementary material, figure S8). This estimate implies a substantial decrease in rates of approximately four orders of magnitude at the tips compared to those at the root.

For lungfishes, 15 PC axes summarised 100% of body shape variance, with 69% captured by the first two axes. PC1 reflects differences in the dorsal extent and anterior reach of the epichordal lobe (52.6% of total variance) and PC2 indicates the anterior extension and separation of the dorsal fin or fins as well as the extent of the anal fin (16.5% of variance) (electronic supplementary material, figure S5). PC3 and PC4 capture two aspects of body depth (extension and compression, respectively; 18% of total variance, cumulatively). Subsequent axes summarised more subtle differences between fin placement, head length, and body length. The best-fit evolutionary model was ACDC (GICw ∼ 0.52), although this is not decisive due to alternative models we evaluated receiving moderate support (Ornstein-Uhlenbeck [OU]: GICw ∼ 0.34; Brownian motion [BM]: GICw ∼ 0.14) (figure 2*c*; electronic supplementary material, table S3). Additionally, some of our results show slightly higher support for OU in fitted posterior trees when ACDC was not the preferred model (figure 2*c*). Estimated *b* was negative (*b* ∼ −3.85; 95% CI from parametric bootstrapping: −24.3, −0.91; electronic supplementary material, figure S8), corresponding to the EB case of ACDC. This value represents an order of magnitude decline in rates, with those at the youngest tips 2% of that at the root. Although the confidence interval for the bootstrapped values does not include zero, some replicates estimated a positive *b*, indicating an inability to distinguish EB from BM (electronic supplementary material, figure S8).

For holosteans, 60 PC axes summarised 100% of body shape variation, with 63% of the variance summarised by the first two axes. PC1 captures the anterior-posterior position of the dorsal fin (35.6% of total variance), and PC2 reflects body depth (27.2% of variance) (electronic supplementary material, figure S6). PC3 and PC4 indicate dorsal and caudal fin form and placement, and dorsal or ventral body curves (together 16.9% of total variance). Further axes summarised differences in snout and head length, fin placement, and body shape. The best-fit evolutionary model was ACDC (GICw ∼ 0.90) (figure 3*c*; electronic supplementary material, table S4). Estimated *b* was weakly negative (*b* ∼ −0.27; 95% CI from parametric bootstrapping: −1.65, 0.44, electronic supplementary material, figure S8), indicating a minor decrease where the rates at the youngest tips are ∼76% of those at the base of the tree. The distribution of b generated by parametric bootstrapping includes zero, which signifies difficulty differentiating between changing versus constant rates (a pattern more readily apparent in the nearly identical plots of GIC scores for holosteans, electronic supplementary material, figure S7).

## 4. Discussion and Conclusions

### (a) Comparison with Previous Interpretations of Phenotypic Evolution in Living Fossil Lineages

Our results provide a reexamination of previous interpretations of phenotypic evolution in coelacanths, lungfishes, and holosteans. While these groups are commonly thought to exhibit specific types of morphological change, the degree to which these evolutionary trajectories have been quantified varies substantially. Patterns in lungfishes are perhaps the most well-established as they benefit from a long pedigree of research, beginning with Westoll’s [17] formative study quantifying the accumulation of “modern” traits in fossil lungfishes. That study and those that followed [14,21,32] support high rates of change in discrete traits at the root of lungfish phylogeny followed by a pronounced decline over the group’s history. Our results broadly corroborate these findings. In contrast to studies of evolutionary rate in discrete characters, there are no quantified assessments of body-shape evolution in lungfishes. Nevertheless, there is a strong informal argument in the literature—tracing back to at least Dollo [89]—that the most substantial changes to lungfish body shape occurred early in the group’s Devonian history, associated with the origin of a single, continuous median fin [32,90,91]. However, we do not find overwhelmingly strong support for declining rates of body shape evolution in lungfishes, with constant-rate alternatives having a combined GICw ∼0.48. The lungfish shape dataset is the smallest for the three lineages considered (*n* = 16, versus 27 coelacanths and 62 holosteans). This sample size is a limitation of the lungfish record [35,89–91] and it is possible additional discoveries of articulated material could increase support for a model with high early rates of change (e.g., EB) for the group.

Quantitative investigation of phenotypic evolution in coelacanths began with Schaeffer’s [33] comparative analysis of the group alongside lungfishes. Inspired by and building on Westoll’s [17] approach, Schaeffer [33] identified similar patterns for lungfishes and coelacanths: a Devonian peak in rates of change followed by a rapid decay. Using a more extensive set of taxa and characters in an explicitly phylogenetic framework, Cloutier [20] reported a broadly similar pattern with three different measures of evolutionary rate. Forey [18] provided a coarse overview of accumulated changes across coelacanth phylogeny between arbitrary stratigraphic intervals, showing a net decline in steps over time. In line with these previous estimates, we find a decrease in rates of discrete character evolution in coelacanths, punctuated by modest increases in the latest Paleozoic and earliest Mesozoic. Interestingly, the most consistent interval of declining rates in coelacanths begins in the early Triassic, preceding their peak in lineage diversity later in that period [18] (electronic supplementary material, figure S1). Thus, it appears that the onset of low rates of change preceded a decline in taxonomic diversity. Whether or not this inference is affected by variability in sampling over time is an unavoidable potential source of bias when examining lineages that become depauperate over time. Nevertheless, our results add further detail to this picture by showing that coelacanths consistently developed new traits even as their overall rates of change declined over time. As with lungfishes, no quantitative assessments of coelacanth evolutionary rates are based on shape. However, Schaeffer ([33]: figure 1) implied extreme conservatism in body form consistent with low rates of change. Friedman and Coates [35] suggested that high morphological disparity early in coelacanth history could stem from elevated evolutionary rates that subsequently declined. Our comparative analyses strongly support this latter interpretation, which indicates a substantial decline in rates of body-shape evolution in coelacanths over their history.

Holosteans are central to the concept of living fossils, but there are few quantitative assessments of phenotypic evolution in the group. Clarke et al. [77] compared patterns of body shape evolution between early holosteans and teleosts, finding no consistent differences in rate between the groups when considering phylogenetic uncertainty. This prior study did not test for variable evolutionary rates within holosteans, and there is no strong prior hypothesis for patterns within the group beyond qualitative assessments based on close phenotypic correspondence between living holosteans relative to their closest fossil relatives. Clarke et al. [77] did investigate the degree of constraint in holostean body shape relative to null models using Bloomberg’s *K*, and suggested that body shape evolution in holosteans is substantially more constrained than that expected under Brownian motion. Clarke and Friedman [32] later reported relatively static levels of morphological disparity over the first 150 million years of holostean history, consistent with myriad modes of phenotypic change, including evolutionary constraint [92]. Although our examination of character saturation for holosteans suggests some degree of constraint for discrete characters, we find that OU is the least well supported of the models considered for holostean shape data.

### (b) Inconsistent Evolutionary Patterns among and within Living Fossil Lineages

Systematic paleobiologists often infer processes or modes of phenotypic evolution from subsets of an organism’s overall anatomy. However, different aspects of morphology can show divergent macroevolutionary patterns, suggesting that mosaic evolution of phenotypes is widespread [30,92]. Therefore, we aimed to summarise the overall anatomy of our focal lineages across two datasets—discrete characters and overall body shape—which has provided an opportunity to examine the degree to which different aspects of phenotype show corresponding or divergent patterns of change. Our unified approach helps to more fully understand the living fossil concept, which has historically been evaluated using independent measures of phenotype. Contrasting approaches and varied anatomical subsets can be found not only in the literature examining our focal clades, but also in groups as taxonomically distant as horseshoe crabs [13] and tuataras [15], where declining disparity and rates over time, respectively, have been recovered for those groups independently.

Two important conclusions emerge from our analyses. First, there does not appear to be a single pattern of morphological evolution universally shared *among* these lineages. Although there is some evidence for declining rates when assessing both discrete-trait and shape evolution separately for all three groups, both the support for, and magnitude of, this decline shows substantial differences among the lineages. Second, and perhaps equally important, different aspects of morphology (i.e., discrete characters and shape) do not imply similar underlying processes *within* each lineage, even though each data type might be thought of as independently, and broadly, summarising overall organismal morphology.

We observe a close correspondence between discrete and continuous trait evolution in coelacanths (Table S1). Rates of discrete-character evolution show a first-order pattern of decline over time; body shape is likewise best fit by a declining-rates ACDC (i.e., EB) model, and parameter estimates imply a reduction of rates by several orders of magnitude over coelacanth history. Patterns of character saturation for coelacanths suggest continued innovation throughout the clade’s history, suggesting few constraints on the evolution of discrete traits (consistent with persistent diffusion through trait space), although at ever-decreasing rates. Likewise, evolutionary patterns across both morphological datasets in holosteans are highly congruent (Table S1). Rates of discrete-character evolution are remarkably constant over time, with only a modest decline over their entire history. Patterns of shape evolution broadly mirror this, with estimated values of *b* suggesting only minor changes in evolutionary rates throughout clade history.

Lungfishes present an alternative case. Like coelacanths, their rates of discrete character evolution decline substantially over time yet, in contrast, lungfishes appear to evolve in a more constrained fashion with more pronounced character saturation. Although the EB case of ACDC is the favoured model for body shape evolution in lungfishes, this is not decisive given its weak support (GICw ∼ 0.52) combined with parameter estimates that imply a modest rate reduction over time in comparison to coelacanths (Table S1).

Phenotypically, lungfishes, coelacanths, and holosteans illustrate attributes commonly ascribed to living fossils [7]: extant species closely resemble some ancient taxa. However, it has long been clear that superficially similar patterns like these can arise from contrasting dynamics of evolutionary change [93]. Phylogenetic information is crucial for distinguishing between alternative mechanisms underlying otherwise comparable phenotypic patterns [94]. By employing a common phylogenetic comparative framework, we sought to determine whether the similarities among three canonical living fossil groups reflected shared aspects of phenotypic evolution or whether contrasting processes yielded superficially similar morphological patterns. We find differences in the support for distinct evolutionary models inferred across—and in some cases within—these living fossil lineages. Thus, as with other quantitative assessments of phenotypic diversification across clades [24], we find that shared features ascribed to living fossils can arise from distinct histories of evolutionary change. We anticipate that a broader approach will lead to better characterization of such groups [29], contributing to a better understanding of the origin and persistence of living fossils and other phylogenetic relics [8].

## Supporting information

Electronic Supplementary Material

## Ethics

This work did not require ethical approval from a human subject or animal welfare committee.

## Data Accessibility

Following acceptance, all code and data will be available in the Dryad Digital Repository and on Github.

## Authors’ contributions

R.A.R.-V.: conceptualization, data curation, formal analysis, investigation, methodology, software, validation, visualisation, writing – original draft, writing – review & editing;

J.S.B: formal analysis, methodology, software, validation, visualisation, writing – original draft, writing – review & editing;

J.T.C.: data curation, validation, writing – original draft, writing – review & editing;

M.F.: conceptualization, investigation, methodology, supervision, validation, writing – original draft, writing – review & editing.

All authors gave final approval for publication and agreed to be held accountable for the work performed herein.

## Declaration of AI use

We have not used AI-assisted technologies in creating this article.

## Acknowledgements

We thank R. Close for providing the initial rate estimation script, J. Clavel for help with discussions on best practices and optimization of the *mvMORPH* model tests, and G. T. Lloyd for assistance with specifics of the character saturation analyses.

## Footnotes

Electronic supplementary material have been uploaded alongside this preprint. Following acceptance, final electronic supplementary materials will be available online at the publisher’s website.

## References

1. Darwin CR. 1859 On the Origin of Species by Means of Natural Selection, or the Preservation of Favoured Races in the Struggle for Life. London: John Murray.

2. Eldredge N, Stanley SM. 1984 Living Fossils. New York: Springer Verlag.

3. Casane D, Laurenti P. 2013 Why coelacanths are not ‘living fossils’. BioEssays 35, 332– 338. (doi:10.1002/bies.201200145)

4. Watkins A. 2021 The Epistemic Value of the Living Fossils Concept. Philos. Sci. 88, 1221– 1233. (doi:10.1086/714875)

5. Grandcolas P, Nattier R, Trewick S. 2014 Relict species: a relict concept? Trends Ecol. Evol. 29, 655–663. (doi:10.1016/j.tree.2014.10.002)

6. Lidgard S, Love AC. 2018 Rethinking Living Fossils. BioScience 68, 760–770. (doi:10.1093/biosci/biy084)

7. Lidgard S, Love AC. 2021 The living fossil concept: reply to Turner. Biol. Philos. 36, 13. (doi:10.1007/s10539-021-09789-z)

8. Turner DD. 2019 In defense of living fossils. Biol. Philos. 34, 23. (doi:10.1007/s10539-019-9678-y)

9. Werth AJ, Shear WA. 2014 The Evolutionary Truth About Living Fossils. Am. Sci. 102, 434– 443.

10. Schopf TJM. 1984 Rates of Evolution and the Notion of ‘Living Fossils’. Annu. Rev. Earth Planet. Sci. 12, 245–292. (doi:10.1146/annurev.ea.12.050184.001333)

11. Fisher DC. 1990 Rates of Evolution – Living Fossils. In Paleobiology: a Synthesis (eds DEG Briggs, PR Crowther), pp. 152–159. London, UK: Blackwell Scientific.

12. Cavin L, Guinot G. 2014 Coelacanths as ‘almost living fossils’. Front. Ecol. Evol. 2, 1–5. (doi:10.3389/fevo.2014.00049)

13. Bicknell RDC et al. 2022 Habitat and developmental constraints drove 330 million years of horseshoe crab evolution. Biol. J. Linn. Soc. 136, 155–172. (doi:10.1093/biolinnean/blab173)

14. Lloyd GT, Wang SC, Brusatte SL. 2011 Identifying heterogeneity in rates of morphological evolution: discrete character change in the evolution of lungfish (Sarcopterygii; Dipnoi). Evolution 66, 330–348. (doi:10.1111/j.1558-5646.2011.01460.x)

15. Simões TR, Caldwell MW, Pierce SE. 2020 Sphenodontian phylogeny and the impact of model choice in Bayesian morphological clock estimates of divergence times and evolutionary rates. BMC Biol. 18, 191. (doi:10.1186/s12915-020-00901-5)

16. Bennett DJ, Sutton MD, Turvey ST. 2018 Quantifying the living fossil concept. *Palaeontol*. Electron. 21. (doi:10.26879/750)

17. Westoll TS. 1949 On the evolution of the Dipnoi. In Genetics, Paleontology and Evolution (eds GL Jepsen, E Mayr, GG Simpson), pp. 121–184. Princeton, NJ: Princeton University Press.

18. Forey PL. 1998 History of the Coelacanth Fishes. Chapman & Hall.

19. Forey PL. 1988 Golden jubilee for the coelacanth *Latimeria chalumnae*. Nat. Rev. 336, 727– 732.

20. Cloutier R. 1991 Patterns, trends, and rates of evolution within the Actinistia. Environ. Biol. Fishes 32, 23–58. (doi:10.1007/BF00007444)

21. Cui X, Friedman M, Qiao T, Yu Y, Zhu M. 2022 The rapid evolution of lungfish durophagy. Nat. Commun. 13, 2390. (doi:10.1038/s41467-022-30091-3)

22. Uyeda JC, Zenil-Ferguson R, Pennell MW. 2018 Rethinking phylogenetic comparative methods. Syst. Biol. 67, 1091–1109. (doi:10.1093/sysbio/syy031)

23. Herrera-Flores JA, Stubbs TL, Benton MJ. 2017 Macroevolutionary patterns in Rhynchocephalia: is the tuatara (*Sphenodon punctatus*) a living fossil? Palaeontology 60, 319–328. (doi:10.1111/pala.12284)

24. Harmon LJ et al. 2010 Early bursts of body size and shape evolution are rare in comparative data. Evolution (doi:10.1111/j.1558-5646.2010.01025.x)

25. Miles DB, Ricklefs RE, Losos JB. 2023 How exceptional are the classic adaptive radiations of passerine birds? Proc. Natl. Acad. Sci. 120, e1813976120. (doi:10.1073/pnas.1813976120)

26. Givnish TJ. 2015 Adaptive radiation versus ‘radiation’ and ‘explosive diversification’: why conceptual distinctions are fundamental to understanding evolution. New Phytol. 207, 297– 303. (doi:10.1111/nph.13482)

27. Osborn HF. 1902 The Law of Adaptive Radiation. Am. Nat. 36, 353–363.

28. Schluter D. 2000 The ecology of adaptive radiation. Oxford: Oxford University Press.

29. Simões M, Breitkreuz L, Alvarado M, Baca S, Cooper JC, Heins L, Herzog K, Lieberman BS. 2016 The Evolving Theory of Evolutionary Radiations. Trends Ecol. Evol. 31, 27–34. (doi:10.1016/j.tree.2015.10.007)

30. Slater GJ, Friscia AR. 2019 Hierarchy in adaptive radiation: A case study using the Carnivora (Mammalia). Evolution 73, 524–539. (doi:10.1111/evo.13689)

31. Musser AM. 2003 Review of the monotreme fossil record and comparison of palaeontological and molecular data. Comp. Biochem. Physiol. A. Mol. Integr. Physiol. 136, 927–942. (doi:10.1016/S1095-6433(03)00275-7)

32. Clarke JT, Friedman M. 2018 Body-shape diversity in Triassic–Early Cretaceous neopterygian fishes: sustained holostean disparity and predominantly gradual increases in teleost phenotypic variety. Paleobiology 44, 402–433. (doi:10.1017/pab.2018.8)

33. Schaeffer B. 1952 Rates of Evolution in the Coelacanth and Dipnoan Fishes. Evolution 6, 101. (doi:10.2307/2405507)

34. Toriño P, Soto M, Perea D. 2021 A comprehensive phylogenetic analysis of coelacanth fishes (Sarcopterygii, Actinistia) with comments on the composition of the Mawsoniidae and Latimeriidae: evaluating old and new methodological challenges and constraints. Hist. Biol. 33, 3423–3443. (doi:10.1080/08912963.2020.1867982)

35. Friedman M, Coates MI. 2006 A newly recognized fossil coelacanth highlights the early morphological diversification of the clade. Proc. R. Soc. B Biol. Sci. 273, 245–250. (doi:10.1098/rspb.2005.3316)

36. Dutel H, Maisey JG, Schwimmer DR, Janvier P, Herbin M, Clément G. 2012 The Giant Cretaceous Coelacanth (Actinistia, Sarcopterygii) *Megalocoelacanthus dobiei* Schwimmer, Stewart & Williams, 1994, and Its Bearing on Latimerioidei Interrelationships. PLoS ONE 7, e49911. (doi:10.1371/journal.pone.0049911)

37. Cavin L, Mennecart B, Obrist C, Costeur L, Furrer H. 2017 Heterochronic evolution explains novel body shape in a Triassic coelacanth from Switzerland. Sci. Rep. 7, 13695. (doi:10.1038/s41598-017-13796-0)

38. Schultze H-P, Chorn J. 1997 The Permo-Carboniferous genus *Sagenodus* and the beginning of modern lungfish. Contrib. Zool. 67, 9–70.

39. Schultze H-P. 2001 *Melanognathus*, a primitive dipnoan from the Lower Devonian of the Canadian arctic and the interrelationships of Devonian dipnoans. J. Vertebr. Paleontol. 21, 781–794. (doi:10.1671/0272-4634(2001)021[0781:MAPDFT]2.0.CO;2)

40. Schultze H-P. 2004 Mesozoic sarcopterygians. In Mesozoic Fishes 3 – Systematics, Paleoenvironments and Biodiversity (eds G Arratia, A Tintori), pp. 463–492. München: Verlag Dr. Friedrich Pfeil.

41. Criswell KE. 2015 The comparative osteology and phylogenetic relationships of African and South American lungfishes (Sarcopterygii: Dipnoi). Zool. J. Linn. Soc. 174, 801–858. (doi:10.1111/zoj.12255)

42. Challands TJ, Smithson TR, Clack JA, Bennett CE, Marshall JEA, Wallace-Johnson SM, Hill H. 2019 A lungfish survivor of the end-Devonian extinction and an Early Carboniferous dipnoan radiation. J. Syst. Palaeontol. 17, 1825–1846. (doi:10.1080/14772019.2019.1572234)

43. Smithson TR, Richards KR, Clack JA. 2016 Lungfish diversity in Romer’s Gap: reaction to the end-Devonian extinction. Palaeontology 59, 29–44. (doi:10.1111/pala.12203)

44. Schultze H-P, Marshall CR. 1993 Contrasting the use of functional complexes and isolated characters in lungfish evolution. Mem. Assoc. Australas. Palaeontol. 15, 211–224.

45. López-Arbarello A, Sferco E. 2018 Neopterygian phylogeny: the merger assay. R. Soc. Open Sci. 5, 172337. (doi:10.1098/rsos.172337)

46. Bouckaert R, Heled J, Kühnert D, Vaughan T, Wu C-H, Xie D, Suchard MA, Rambaut A, Drummond AJ. 2014 BEAST 2: A Software Platform for Bayesian Evolutionary Analysis. PLoS Comput. Biol. 10, e1003537. (doi:10.1371/journal.pcbi.1003537)

47. Bouckaert R et al. 2019 BEAST 2.5: An advanced software platform for Bayesian evolutionary analysis. PLOS Comput. Biol. 15, e1006650. (doi:10.1371/journal.pcbi.1006650)

48. Stadler T. 2010 Sampling-through-time in birth–death trees. J. Theor. Biol. 267, 396–404. (doi:10.1016/j.jtbi.2010.09.010)

49. Heath TA, Huelsenbeck JP, Stadler T. 2014 The fossilized birth-death process for coherent calibration of divergence-time estimates. Proc. Natl. Acad. Sci. 111, E2957–E2966. (doi:10.1073/pnas.1319091111)

50. Arcila D, Pyron RA, Tyler JC, Ortí G, Betancur-R R. 2015 An evaluation of fossil tip-dating versus node-age calibrations in tetraodontiform fishes (Teleostei: Percomorphaceae). Mol. Phylogenet. Evol. 82, 131–145. (doi:10.1016/j.ympev.2014.10.011)

51. Grimm GW, Kapli P, Bomfleur B, McLoughlin S, Renner SS. 2015 Using More Than the Oldest Fossils: Dating Osmundaceae with Three Bayesian Clock Approaches. Syst. Biol. 64, 396–405. (doi:10.1093/sysbio/syu108)

52. Zhang C, Stadler T, Klopfstein S, Heath TA, Ronquist F. 2016 Total-Evidence Dating under the Fossilized Birth–Death Process. Syst. Biol. 65, 228–249. (doi:10.1093/sysbio/syv080)

53. Luo A, Duchêne DA, Zhang C, Zhu C-D, Ho SYW. 2019 A Simulation-Based Evaluation of Tip-Dating Under the Fossilized Birth–Death Process. Syst. Biol. **syz**038. (doi:10.1093/sysbio/syz038)

54. Lewis PO. 2001 A likelihood approach to estimating phylogeny from discrete morphological character data. Syst. Biol. 50, 913–925. (doi:10.1080/106351501753462876)

55. Zhu M, Zhao W, Jia L, Lu J, Qiao T, Qu Q. 2009 The oldest articulated osteichthyan reveals mosaic gnathostome characters. Nature 458, 469–474. (doi:10.1038/nature07855)

56. Matzke NJ, Wright A. 2016 Inferring node dates from tip dates in fossil Canidae: the importance of tree priors. Biol. Lett. 12, 20160328. (doi:10.1098/rsbl.2016.0328)

57. Kemp A, Cavin L, Guinot G. 2017 Evolutionary history of lungfishes with a new phylogeny of post-Devonian genera. Palaeogeogr. Palaeoclimatol. Palaeoecol. 471, 209–219. (doi:10.1016/j.palaeo.2016.12.051)

58. Longrich NR. 2017 A stem lepidosireniform lungfish (Sarcopterygia: Dipnoi) from the Upper Eocene of Libya, North Africa and implications for Cenozoic lungfish evolution. Gondwana Res. 42, 140–150. (doi:10.1016/j.gr.2016.09.007)

59. Capobianco A, Friedman M. 2019 Vicariance and dispersal in southern hemisphere freshwater fish clades: a palaeontological perspective. Biol. Rev. 94, 662–699. (doi:10.1111/brv.12473)

60. Grande L. 2010 An Empirical and Synthetic Pattern Study of Gars (Lepisosteiformes) and Closely Related Species, Based Mostly on Skeletal Anatomy. The Resurrection of Holostei. Copeia 10, 1–871.

61. Rambaut A, Drummond AJ, Xie D, Baele G, Suchard MA. 2018 Posterior Summarization in Bayesian Phylogenetics Using Tracer 1.7. Syst. Biol. 67, 901–904. (doi:10.1093/sysbio/syy032)

62. Gradstein FM, Ogg JG, Schmitz MD, Ogg GM. 2012 The Geologic Time Scale 2012. 1st ed. Amsterdam; Boston: Elsevier.

63. Ogg JG, Ogg GM, Gradstein FM. 2016 A Concise Geologic Time Scale 2016. Amsterdam, Netherlands: Elsevier.

64. Gradstein FM, Ogg JG, Schmitz MD, Ogg GM. 2020 The Geologic Time Scale 2020. Amsterdam: Elsevier.

65. King B, Rücklin M. 2020 Tip dating with fossil sites and stratigraphic sequences. PeerJ 8, e9368. (doi:10.7717/peerj.9368)

66. Revell LJ. 2012 phytools: an R package for phylogenetic comparative biology (and other things). Methods Ecol. Evol. 3, 217–223. (doi:10.1111/j.2041-210X.2011.00169.x)

67. Paradis E, Claude J, Strimmer K. 2004 APE: Analyses of Phylogenetics and Evolution in R language. Bioinformatics 20, 289–290. (doi:10.1093/bioinformatics/btg412)

68. Plummer M, Best N, Cowles K, Vines K. 2006 CODA: Convergence diagnosis and output analysis for MCMC. R News 6, 7–11.

69. Close RA, Johanson Z, Tyler JC, Harrington RC, Friedman M. 2016 Mosaicism in a new Eocene pufferfish highlights rapid morphological innovation near the origin of crown tetraodontiforms. Palaeontology 59, 499–514. (doi:10.1111/pala.12245)

70. Jombart T et al. 2014 OutbreakTools: A new platform for disease outbreak analysis using the R software. Epidemics 7, 28–34. (doi:10.1016/j.epidem.2014.04.003)

71. R Core Team. 2013 R: A language and environment for statistical computing.

72. Wagner PJ. 2000 Exhaustion of morphologic character states among fossil taxa. Evolution 54, 365–386. (doi:10.1111/j.0014-3820.2000.tb00040.x)

73. Wagner PJ, Ruta M, Coates MI. 2006 Evolutionary patterns in early tetrapods. II. Differing constraints on available character space among clades. Proc. R. Soc. B Biol. Sci. 273, 2113–2118. (doi:10.1098/rspb.2006.3561)

74. Oyston JW, Hughes M, Wagner PJ, Gerber S, Wills MA. 2015 What limits the morphological disparity of clades? Interface Focus 5, 20150042. (doi:10.1098/rsfs.2015.0042)

75. Brocklehurst N, Benson RJ. 2021 Multiple paths to morphological diversification during the origin of amniotes. *Nat*. Ecol. Evol. 5, 1243–1249. (doi:10.1038/s41559-021-01516-x)

76. Brocklehurst N, Panciroli E, Benevento GL, Benson RBJ. 2021 Mammaliaform extinctions as a driver of the morphological radiation of Cenozoic mammals. Curr. Biol. 31, 2955–2963.e4. (doi:10.1016/j.cub.2021.04.044)

77. Clarke JT, Lloyd GT, Friedman M. 2016 Little evidence for enhanced phenotypic evolution in early teleosts relative to their living fossil sister group. Proc. Natl. Acad. Sci. 113, 11531– 11536. (doi:10.1073/pnas.1607237113)

78. Olsen AM, Westneat MW. 2015 StereoMorph: an R package for the collection of 3D landmarks and curves using a stereo camera set-up. Methods Ecol. Evol. 6, 351–356. (doi:10.1111/2041-210X.12326)

79. Adams DC, Otárola-Castillo E. 2013 geomorph: an r package for the collection and analysis of geometric morphometric shape data. Methods Ecol. Evol. 4, 393–399. (doi:10.1111/2041-210X.12035)

80. Baken EK, Collyer ML, Kaliontzopoulou A, Adams DC. 2021 geomorph v4.0 and gmShiny: Enhanced analytics and a new graphical interface for a comprehensive morphometric experience. Methods Ecol. Evol. 12, 2355–2363. (doi:10.1111/2041-210X.13723)

81. Clavel J, Escarguel G, Merceron G. 2015 mvMORPH: an R package for fitting multivariate evolutionary models to morphometric data. Methods Ecol. Evol. 6, 1311–1319. (doi:10.1111/2041-210X.12420)

82. Clavel J, Aristide L, Morlon H. 2019 A Penalized Likelihood Framework for High-Dimensional Phylogenetic Comparative Methods and an Application to New-World Monkeys Brain Evolution. Syst. Biol. 68, 93–116. (doi:10.1093/sysbio/syy045)

83. Hansen TF. 1997 Stabilizing selection and the comparative analysis of adaptation. Evolution 51, 1341–1351. (doi:10.1111/j.1558-5646.1997.tb01457.x)

84. Ho LST, Ané C. 2014 Intrinsic inference difficulties for trait evolution with Ornstein-Uhlenbeck models. Methods Ecol. Evol. 5, 1133–1146. (doi:10.1111/2041-210X.12285)

85. Blomberg SP, Garland Jr. T, Ives AR. 2003 Testing for phylogenetic signal in comparative data: behavioral traits are more labile. Evolution 57, 717–745. (doi:10.1111/j.0014-3820.2003.tb00285.x)

86. Silvestro D, Kostikova A, Litsios G, Pearman PB, Salamin N. 2015 Measurement errors should always be incorporated in phylogenetic comparative analysis. Methods Ecol. Evol. 6, 340–346. (doi:10.1111/2041-210X.12337)

87. Konishi S, Kitagawa, Genshiro. 1996 Generalised information criteria in model selection. Biometrika 83, 875–890. (doi:10.1093/biomet/83.4.875)

88. Konishi S, Kitagawa G. 2008 *Information Criteria and Statistical Modeling*. New York, NY: Springer New York. (doi:10.1007/978-0-387-71887-3)

89. Dollo L. 1895 Sur la phylogénie des Dipneustes. Bull. Société Belge Géologie Paléontol. Hydrol. 9, 79–128.

90. Ahlberg PE, Trewin NH. 1994 The postcranial skeleton of the Middle Devonian lungfish *Dipterus valenciennesi*. Trans. R. Soc. Edinb. Earth Sci. 85, 159–175. (doi:10.1017/S0263593300003588)

91. Friedman M, Daeschler EB. 2006 Late Devonian (Famennian) lungfishes from the Catskill Formation of Pennsylvania, USA. Palaeontology 49, 1167–1183. (doi:10.1111/j.1475-4983.2006.00594.x)

92. Hopkins MJ, Lidgard S. 2012 Evolutionary mode routinely varies among morphological traits within fossil species lineages. Proc. Natl. Acad. Sci. 109, 20520–20525. (doi:10.1073/pnas.1209901109)

93. Foote M. 1996 Models of morphological diversification. In Evolutionary Paleobiology (eds D Jablonski, DH Erwin, JH Lipps), pp. 62–86. Chicago: University of Chicago Press.

94. Sidlauskas B. 2008 Continuous and arrested morphological diversification in sister clades of characiform fishes: a phylomorphospace approach. Evolution 62, 3135–3156. (doi:10.1111/j.1558-5646.2008.00519.x)

