## Supplementary material for "Variable patterns of phenotypic evolution among canonical ‘living fossil’ lineages": Electronic Supplementary Material

This is the electronic supplementary material for Rivero-Vega et al. (XXXX) *Variable patterns of phenotypic evolution among canonical 'living fossil' lineages*.

#### S1. Phylogenetic Reconstruction

(a) Coelacanths

The Devonian *Gavinia* + *Styloichthys* were the earliest diverging actinistian clade (90.4% posterior support), followed by *Miguashaia* (45.4%) (figure S1). This is opposite to the arrangement found by Toriño et al. [1], though the earliest portion of their tree had lower bootstrap values than our posterior support (17% – 22%). Our constrained monophyly of *Allenypterus* + *Holopterygius* resulted in *Euporosteus* + *Diplocercides* forming a low-support clade (13%), as opposed to a polytomy. The Carboniferous backbone of our tree, including *Lochmocercus*, *Hadronector*, *Polysteorhynchus*, *Caridosuctor*, and *Rhabdoderma*, was recovered in a different arrangement but with higher support (60% – 92.6%) than Toriño et al. [1]. The Permian lineages also differed in arrangement to those found by Toriño et al. [1], but with comparably low levels of support, which has the hallmarks of a rapid diversification event. Apart from causing issues when inferring relationships, this uncertainty has been identified by previous studies, which have noted its effects alongside issues caused by taxa that have been historically difficult to code [1,2]. *Heptanema* and *Dobrogeria* were recovered within Latimerioidei, with *Whiteia* as sister group to the clade. Within Latimerioidei, the mawsoniid and latimerid relationships had much higher support than in Toriño et al. ([1]: figures 3, 5) (figure S1). *Chinela* and *Diplurus*were were aligned with the mawsoniids and *Garnbergia* was the immediate sister lineage to the latimerids. The earliest diverging latimeriid clade was *Foreyia* + *Ticinepomis* (97.5%), with *Libys* + *Megalocoelacanthus* (87.9% posterior support) found closer to the crown, a reversal of the relationships recovered in Toriño et al. [1]. The Late Cretaceous *Macropoma* is the immediate sister lineage of the extant *Latimeria* (70.5%), with Late Jurassic *Swenzia* the closest relative to this pair, reflecting one [3] of two [1] competing hypotheses of relationships among these lineages. Given the overall higher support coupled with the stratigraphic uncertainty included in our analyses, these results are the most consistent and rigorous for this clade.

(b) Lungfishes

There was extremely low support within and between the majority of Devonian and Permian clades (figure S2), representing the majority of taxa. This resulted in broad differences between the trees estimated in our analyses, and those in Lloyd et al. [4]. Most of the sister-group relationships that had the highest support (>50% posterior support) in our analyses were also found in the original publication (e.g., *Dipnorhynchus spp.*, *Barwickia downunda* + *Dipterus valenciennesi*, *Rhinodipterus spp.*, *Holodipterus spp.*, and *Griphognathus spp.*); however, some relationships with high support differed (e.g., *Palaeodaphus insignis* + [*Jarvickia arctica* + *Sunwapta grandiceps*], [*Rhynchodipterus elginensis* + *Soederberghia groenlandica*] + *Griphognathus spp.*, and *Delatitia breviceps* + *Parasagenodus sibiricus*). The highest backbone support was recovered between the Carboniferous and post-Permian clades (>50%). All these nodes differed from Lloyd et al. [4] except for *Arganodus atlantis* + *Ferganoceratodus jurassicus* (51%). This low support result is likely representative of the quality and construction of these lungfish character datasets over time, as has been noted in prior publications (e.g., [5]).

(c) Holosteans

The topology of the tree was broadly consistent with that in López-Arbarello and Sferco [6], including modest overall support across the tree (figure S3). Here, we will only discuss differences as they pertain to holosteans to the exclusion of any outgroups. Although the three species of *Dapedium* were recovered as stem Holostei on the MCC tree in agreement with recent studies [6,7], they had <50% posterior support, and shorter runs frequently recovered them within Ginglymodi. Because of this, we cannot rule out the group being stem Ginglymodi [8,9] nor any other placement about the holostean stem (e.g., [10]). They are therefore considered early branching holosteans with uncertain phylogenetic placement. Other differences from López-Arbarello and Sferco [6] included *Caturus furcatus* recovered as sister to the Amiiformes rather than to the ionoscopids (58.2%), *Kyphosichthys grandei* nested within *Sangiorgioichthys* (83%), and better resolution within the Semionotiformes and Cretaceous and Paleogene Lepisosteiformes (66% – 100%).

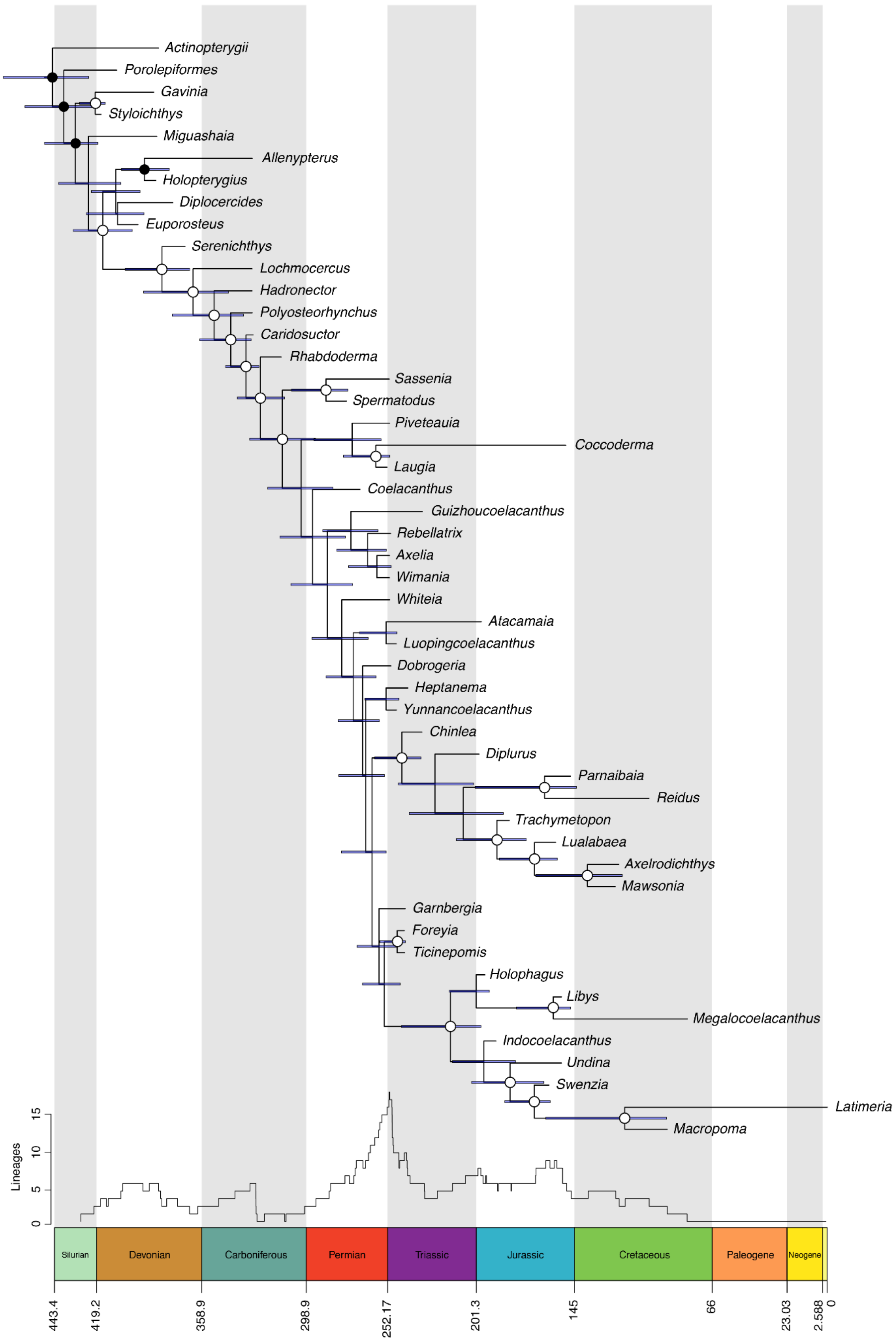

Actinopterygii
Porolepiformes
Gavinia
Styloichthys
Miguashaia
Allenypterus
Holopterygius
Diplocercides
Euporosteus
Serenichthys
Lochmocercus
Hadronector
Polyosteorhynchus
Caridosuctor
Rhabdoderma
Sassenia
Spermatodus
Piveteauia
Coccoderma
Laugia
Coelacanthus
Guizhoucoelacanthus
Rebellatrix
Axelia
Wimania
Whiteia
Atacamaia
Luopingcoelacanthus
Dobrogeria
Heptanema
Yunnancoelacanthus
Chinlea
Diplurus
Parnaibaia
Reidus
Trachymetopon
Lualabaea
Axelrodichthys
Mawsonia
Garnbergia
Foreyia
Ticinepomis
Holophagus
Libys
Megalocoelacanthus
Indocoelacanthus
Undina
Swenzia
Latimeria
Macropoma
Lineages
0
5
10
15
Silurian
Devonian
Carboniferous
Permian
Triassic
Jurassic
Cretaceous
Paleogene
Neogene
443.4
419.2
358.9
298.9
252.17
201.3
145
66
23.03
2.588
0

**Figure S1.** Time-scaled maximum clade credibility tree for coelacanths. Phylogeny inferred using morphological data in BEAST v2.6.5 with the Fossilized Birth-Death model. Nodes marked in black represent clade constraints, nodes marked in black circling white represent clades with >50% posterior probability, and blue bars on nodes represent 95% posterior probability distribution. The graph above the timescale is a Lineages Through Time plot of taxa belonging to the coelacanth clade calculated from the MCC tree.

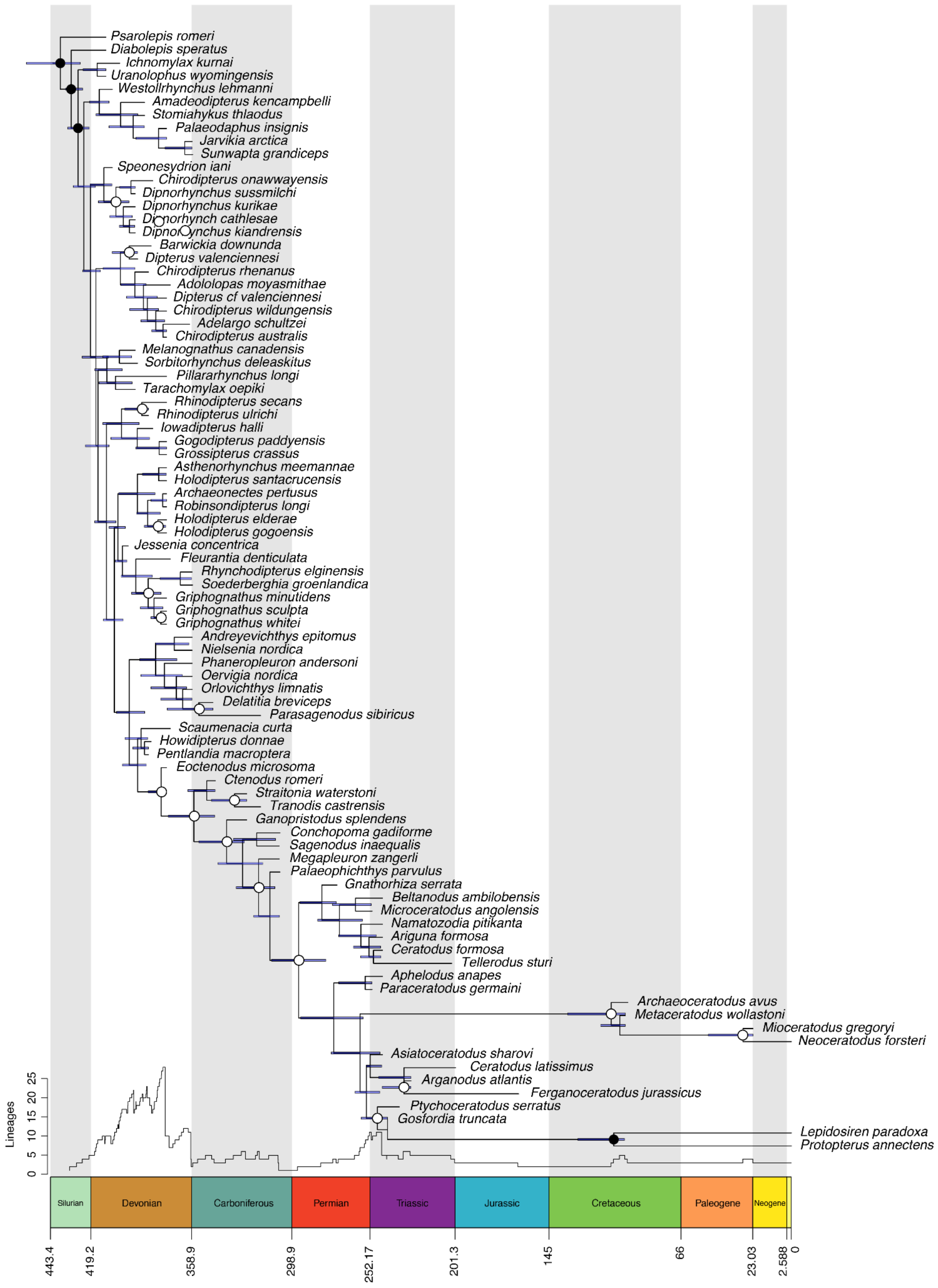

Psarolepis romeri
Diabolepis speratus
Ichnomylax kurnai
Uranolophus wyomingensis
Westollrhynchus lehmanni
Amadeodipterus kencampbelli
Stomiahykus thlaodus
Palaeodaphus insignis
Jarvikia arctica
Sunwapta grandiceps
Speonesydrion iani
Chirodipterus onawwayensis
Dipnorhynchus sussmilchi
Dipnorhynchus kurikae
Dipnorhynch cathlesae
Dipnorhynchus kiandrensis
Barwickia downunda
Dipterus valenciennesi
Chirodipterus rhenanus
Adololopas moyasmithae
Dipterus cf valenciennesi
Chirodipterus wildungensis
Adelargo schultzei
Chirodipterus australis
Melanognathus canadensis
Sorbitorhynchus deleaskitus
Pillararhynchus longi
Tarachomylax oepiki
Rhinodipterus secans
Rhinodipterus ulrichi
Iowadipterus halli
Gogodipterus paddyensis
Grossipterus crassus
Asthenorhynchus meemannae
Holodipterus santacrucensis
Archaeonectes pertusus
Robinsondipterus longi
Holodipterus elderae
Holodipterus gogoensis
Jessenia concentrica
Fleurantia denticulata
Rhynchodipterus elginensis
Soederberghia groenlandica
Griphognathus minutidens
Griphognathus sculpta
Griphognathus whitei
Andreyevichthys epitomus
Nielsenia nordica
Phaneropleuron andersoni
Oervigia nordica
Orlovichthys limnatis
Delatitia breviceps
Parasagenodus sibiricus
Scaumenacia curta
Howidipterus donnae
Pentlandia macroptera
Eoctenodus microsoma
Ctenodus romeri
Straitonia waterstoni
Tranodis castrensis
Ganopristodus splendens
Conchopoma gadiforme
Sagenodus inaequalis
Megapleuron zangerli
Palaeophichthys parvulus
Gnathorhiza serrata
Beltanodus ambilobensis
Microceratodus angolensis
Namatozodia pitikanta
Ariguna formosa
Ceratodus formosa
Tellerodus sturi
Aphelodus anapes
Paraceratodus germaini
Archaeoceratodus avus
Metaceratodus wollastoni
Mioceratodus gregoryi
Neoceratodus forsteri
Asiatoceratodus sharovi
Ceratodus latissimus
Arganodus atlantis
Ferganoceratodus jurassicus
Ptychoceratodus serratus
Gosfordia truncata
Lepidosiren paradoxa
Protopterus annectens
Lineages
0
5
10
15
20
25
Silurian
Devonian
Carboniferous
Permian
Triassic
Jurassic
Cretaceous
Paleogene
Neogene
443.4
419.2
358.9
298.9
252.17
201.3
145
66
23.03
2.588
0

**Figure S2.** Time-scaled maximum clade credibility tree for lungfishes. Phylogeny inferred using morphological data in BEAST v2.6.5 with the Fossilized Birth-Death model. Nodes marked in black represent clade constraints, nodes marked in black circling white represent clades with >50% posterior probability, and blue bars on nodes represent 95% posterior probability distribution. The graph above the timescale is a Lineages Through Time plot of taxa belonging to the lungfish clade calculated from the MCC tree.

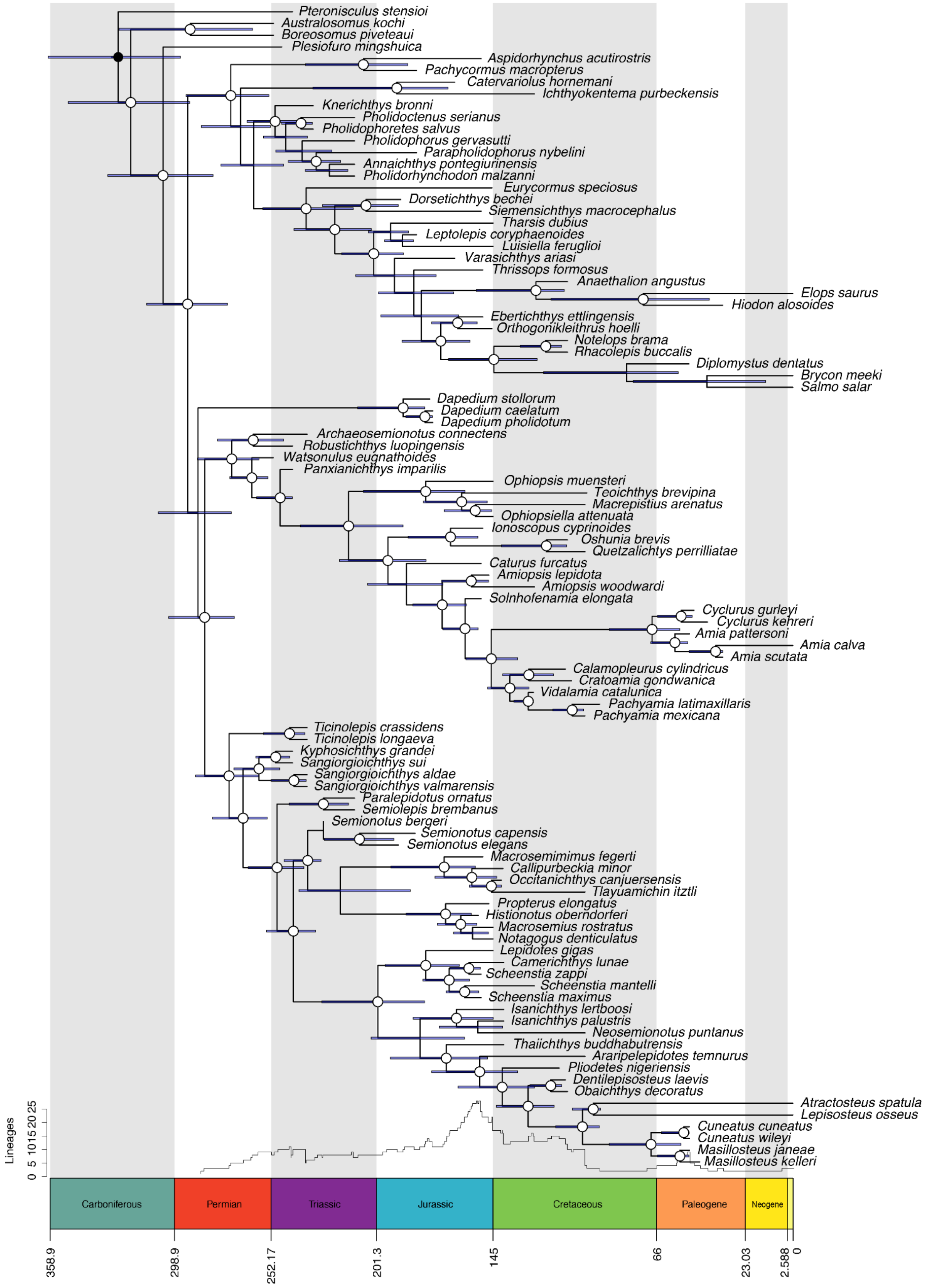

Pteronisculus stensioi
Australosomus kochi
Boreosomus piveteaui
Plesiofuro mingshuica
Aspidorhynchus acutirostris
Pachycormus macropterus
Catervariolus hornemani
Ichthyokentema purbeckensis
Knerichthys bronni
Pholidoctenus serianus
Pholidophoretes salvus
Pholidophorus gervasutti
Parapholidophorus nybelini
Annaichthys pontegiurinensis
Pholidorhynchodon malzanni
Eurycormus speciosus
Dorsetichthys bechei
Siemensichthys macrocephalus
Tharsis dubius
Leptolepis coryphaenoides
Luisiella feruglioi
Varasichthys ariasi
Thrissops formosus
Anaethalion angustus
Elops saurus
Hiodon alosoides
Ebertichthys ettlingensis
Orthogonikleithrus hoelli
Notelops brama
Rhacolepis buccalis
Diplomystus dentatus
Brycon meeki
Salmo salar
Dapedium stollorum
Dapedium caelatum
Dapedium pholidotum
Archaeosemionotus connectens
Robustichthys luopingensis
Watsonulus eugnathoides
Panxianichthys imparilis
Ophiopsis muensteri
Teoichthys brevipina
Macrepistius arenatus
Ophiopsiella attenuata
Ionoscopus cyprinoides
Oshunia brevis
Quetzalichtys perrilliatae
Caturus furcatus
Amiopsis lepidota
Amiopsis woodwardi
Solnhofenamia elongata
Cyclurus gurleyi
Cyclurus kehreri
Amia pattersoni
Amia calva
Amia scutata
Calamopleurus cylindricus
Cratoamia gondwanica
Vidalamia catalunica
Pachyamia latimaxillaris
Pachyamia mexicana
Ticinolepis crassidens
Ticinolepis longaeva
Kyphosichthys grandei
Sangiorgioichthys sui
Sangiorgioichthys aldae
Sangiorgioichthys valmarensis
Paralepidotus ornatus
Semiolepis brembanus
Semionotus bergeri
Semionotus capensis
Semionotus elegans
Macrosemimimus fegerti
Callipurbeckia minor
Occitanichthys canjuersensis
Tlayuamichin itztli
Propterus elongatus
Histionotus oberndorferi
Macrosemius rostratus
Notagogus denticulatus
Lepidotes gigas
Camerichthys lunae
Scheenstia zappi
Scheenstia mantelli
Scheenstia maximus
Isanichthys lertboosi
Isanichthys palustris
Neosemionotus puntanus
Thaiichthys buddhabutrensis
Araripelepidotes temnurus
Pliodetes nigeriensis
Dentilepisosteus laevis
Obaichthys decoratus
Atractosteus spatula
Lepisosteus osseus
Cuneatus cuneatus
Cuneatus wileyi
Masillosteus janeae
Masillosteus kelleri
Lineages
0 5 10 15 20 25
Carboniferous
Permian
Triassic
Jurassic
Cretaceous
Paleogene
Neogene
358.9
298.9
252.17
201.3
145
66
23.03
2.588
0

**Figure S3.** Time-scaled maximum clade credibility tree for holosteans. Phylogeny inferred using morphological data in BEAST v2.6.5 with the Fossilized Birth-Death model. Nodes marked in black represent clade constraints, nodes marked in black circling white represent clades with >50% posterior probability, and blue bars on nodes represent 95% posterior probability distribution. The graph above the timescale is a Lineages Through Time plot of taxa belonging to the holostean clade calculated from the MCC tree.

**S2. Landmarking & Shape Data**

(a) Coelacanths

11 fixed landmarks for coelacanths from Friedman and Coates [11] (1-10, 12) with 3 added in this analysis (11, 13-14) (figure S4): (1) tip of snout, (2) posterior margin of postparietals, (3) anterior insertion of first dorsal fin, (4) posterior insertion of first dorsal fin, (5) posterior insertion of second dorsal fin, (6) anterior insertion of epichordal lobe, (7) posterior tip of accessory lobe, (8) anterior insertion of hypochordal lobe, (9) posterior insertion of anal fin, (10) base of pelvic fin, (11) posterior ventral edge of interpalate, (12) quadrate/articular joint, (13) central, ventral surface of orbit, and (14) central, dorsal surface of orbit.

(b) Lungfishes

11 landmarks were created for lungfishes in this analysis (figure S5): (1) anterior tip of the upper jaw (premaxilla), (2) posteriormost edge of skull roof (posterior to A bone), (3) anterior insertion of dorsal fin, (4) posterior tip of epichordal lobe, (5) anterior insertion of anal fin, (6) anterior insertion of pelvic fin, (7) anterior insertion of the pectoral fin, (8) posterior ventral edge of interpalate, (9) lower jaw joint, (10) central, ventral surface of the orbit, and (11) central, dorsal surface of the orbit. These landmarks were specifically tailored to complement the ones derived from previous analyses for coelacanths and holosteans.

(c) Holosteans

12 fixed landmarks for holosteans from Clarke and Friedman [12] (1-8, 10, 12-14) with two added in this analysis (9, 11) (figure S6): (1) anterior tip of the upper jaw (premaxilla), (2) postero-dorsal tip of braincase, (3) anterior insertion of dorsal fin, (4) posterior insertion of

dorsal fin, (5) dorsal surface representation of the last vertebral centra, (6) ventral surface representation of the last vertebral centra, (7) posterior insertion of anal fin, (8) anterior insertion of anal fin, (9) base of pelvic fin, (10) anterior insertion of the pectoral fin, (11) posterior ventral edge of interpalate, (12) lower jaw, joint, (13) the central, ventral surface of the orbit, and (14) the central, dorsal surface of the orbit. Landmark #2 from Clarke and Friedman [12] is not included in this analysis.

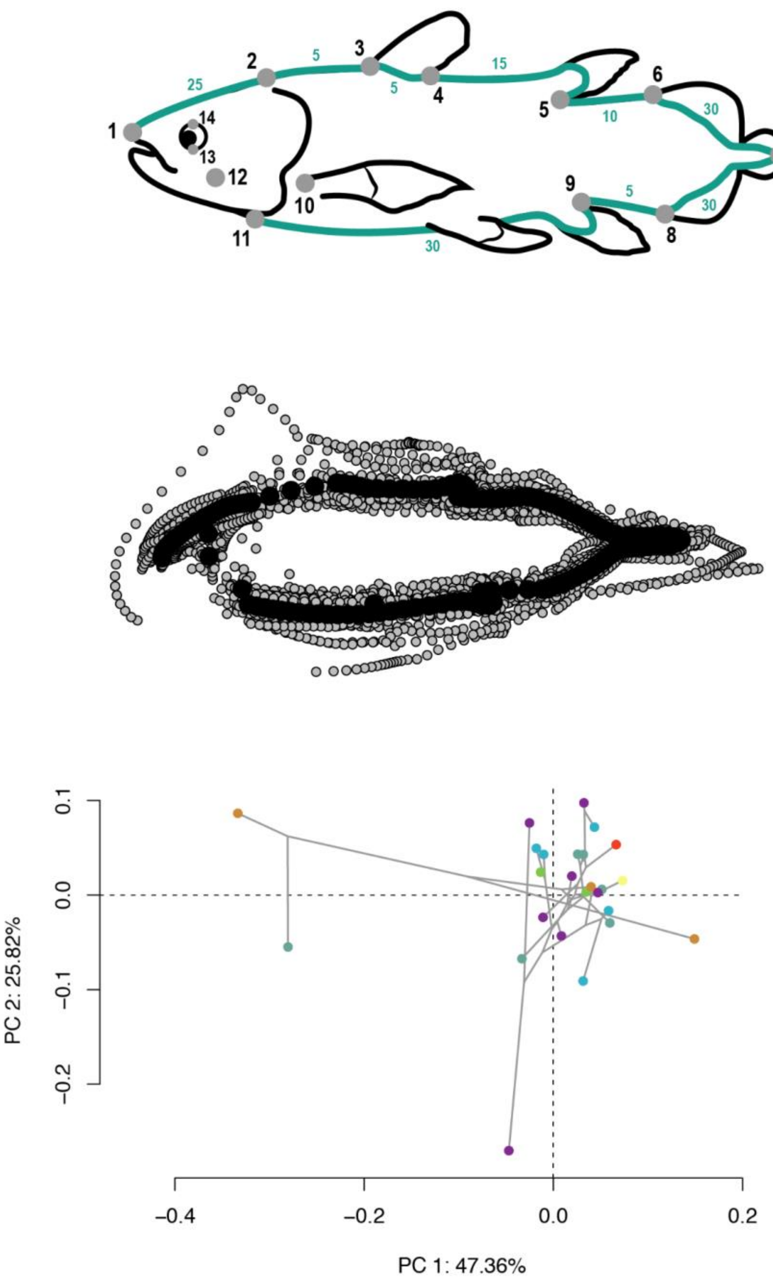


**Figure S4.** Landmarking scheme for coelacanths. The landmarking scheme (top) used to create coordinate data (centre) for morphometric analyses (bottom). The scheme is described in detail in the Landmarking & Shape Data section. Colours representing the last appearance datum of each taxon standardised to the International Commission on Stratigraphy.

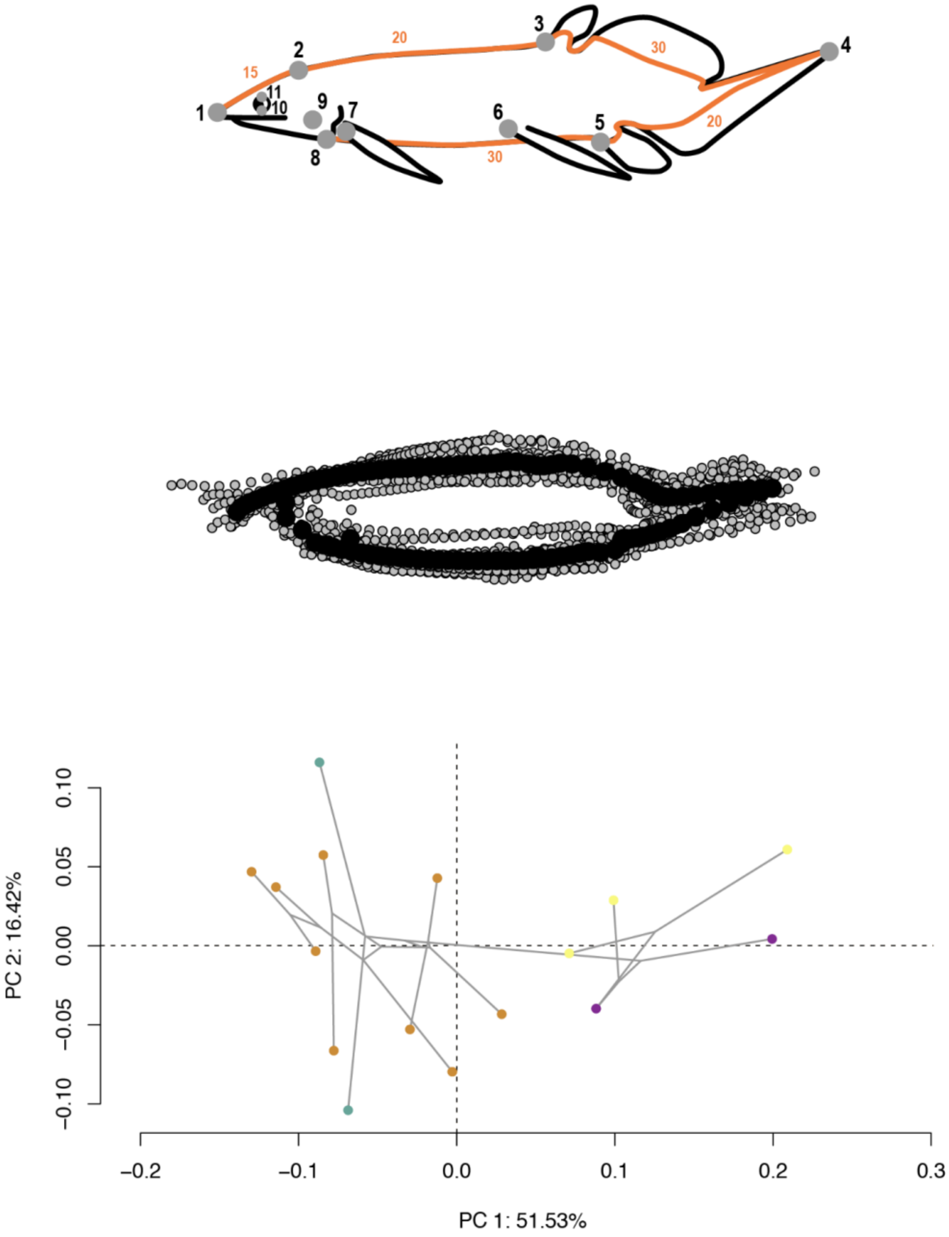


**Figure S5.** Landmarking scheme for lungfishes. The landmarking scheme (top) used to create coordinate data (centre) for morphometric analyses (bottom). The scheme is described in detail in the Landmarking & Shape Data section. Colours representing the last appearance datum of each taxon standardised to the International Commission on Stratigraphy.

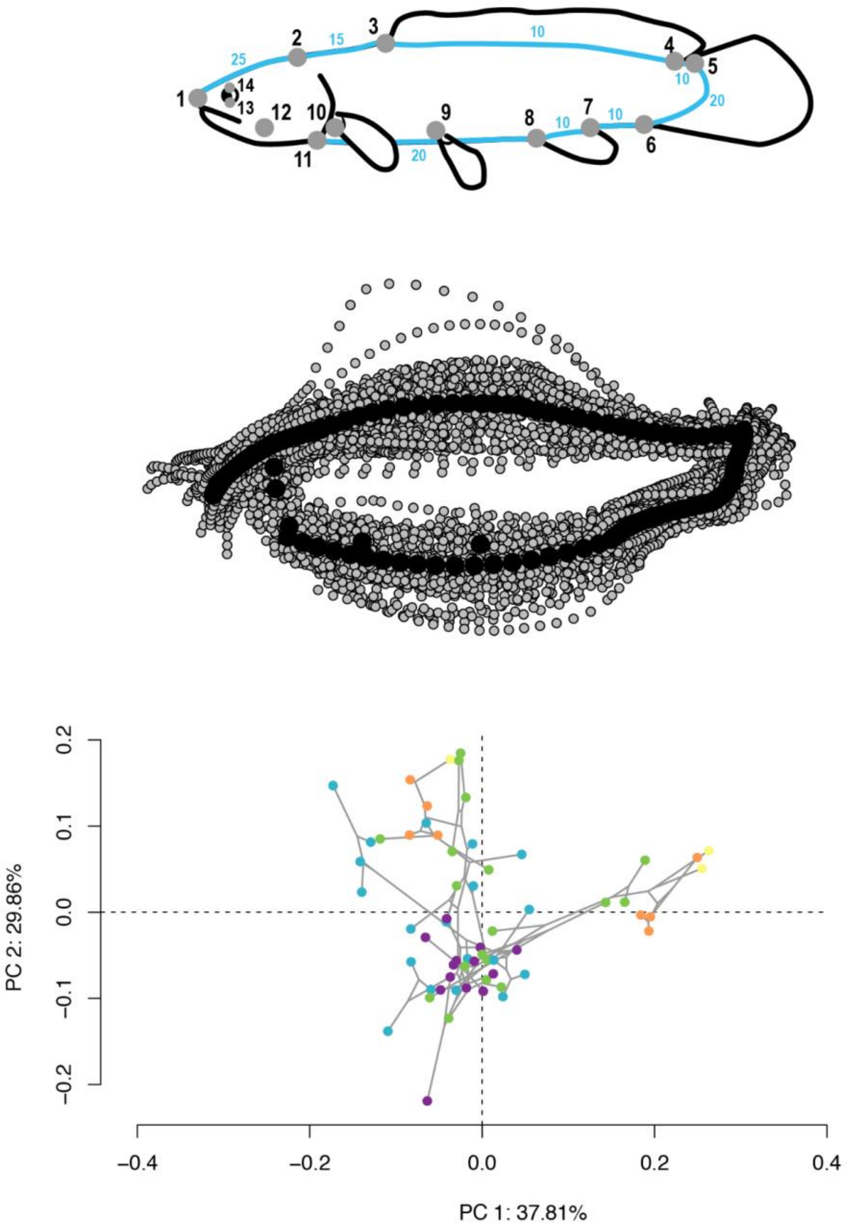


**Figure S6.** Landmarking scheme for holosteans. The landmarking scheme (top) used to create coordinate data (centre) for morphometric analyses (bottom). The scheme is described in detail in the Landmarking & Shape Data section. Colours representing the last appearance datum of each taxon standardised to the International Commission on Stratigraphy.

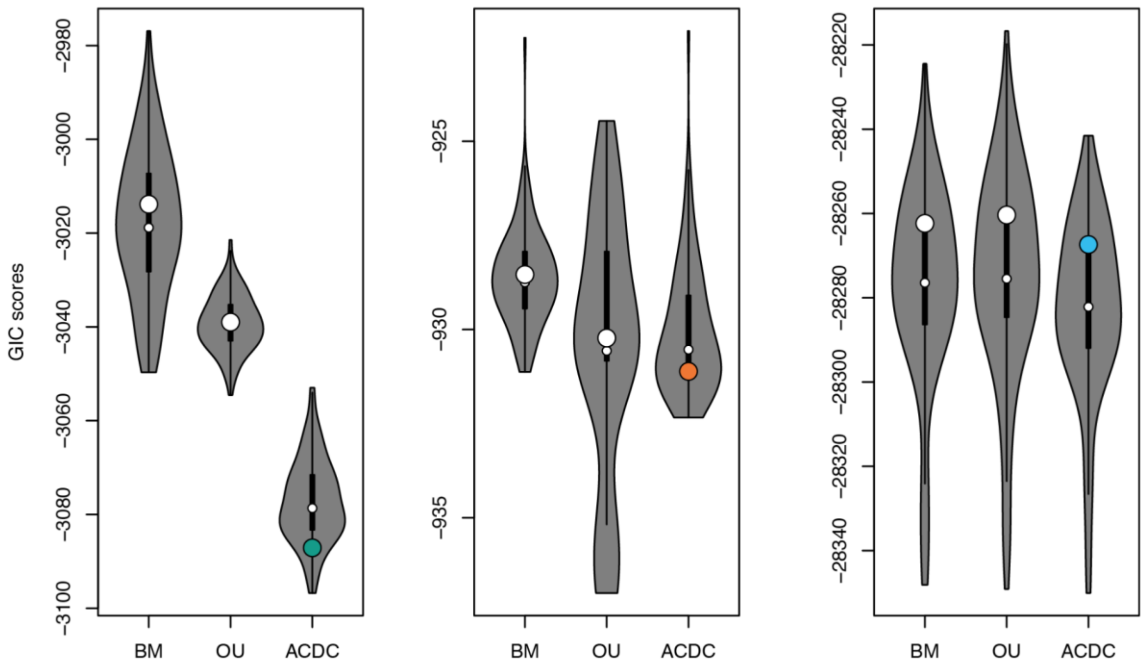


**Figure S7.** GIC scores for each group. We calculated GIC scores for coelacanths (left), lungfishes (centre), and holosteans (right). Model fitting was performed via *mvMORPH* and the PC axes that summarised 100% of the variability for each clade using the MCC tree (circles) and 100 trees randomly sampled from the posterior (violin plot distribution).

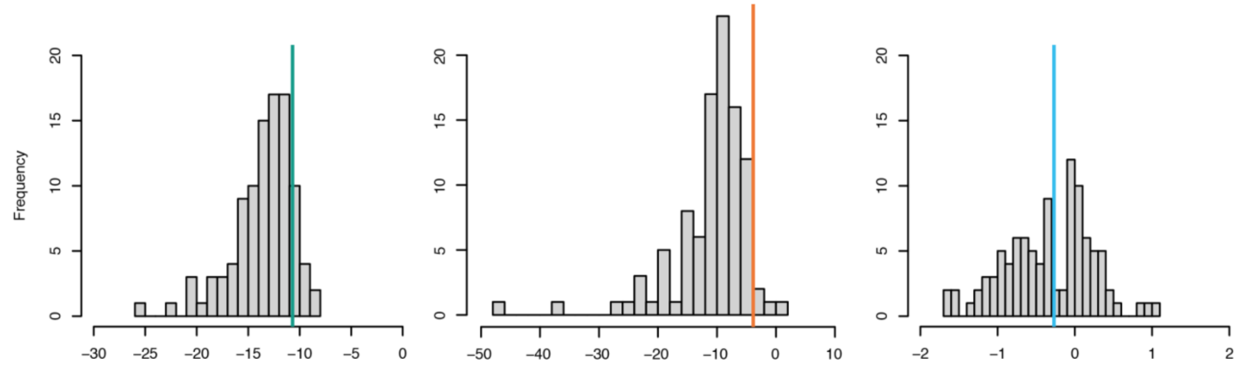


**Figure S8.** Bootstrapped beta parameter estimates for each group. We performed 100 bootstrap replicates for coelacanths (left), lungfishes (centre), and holosteans (right) using the estimates from the empirical model fitting. Vertical coloured bars indicate the MCC-derived parameter estimate for each group (-10.7, -3.85, and -0.27 for coelacanths, lungfishes, and holosteans, respectively).

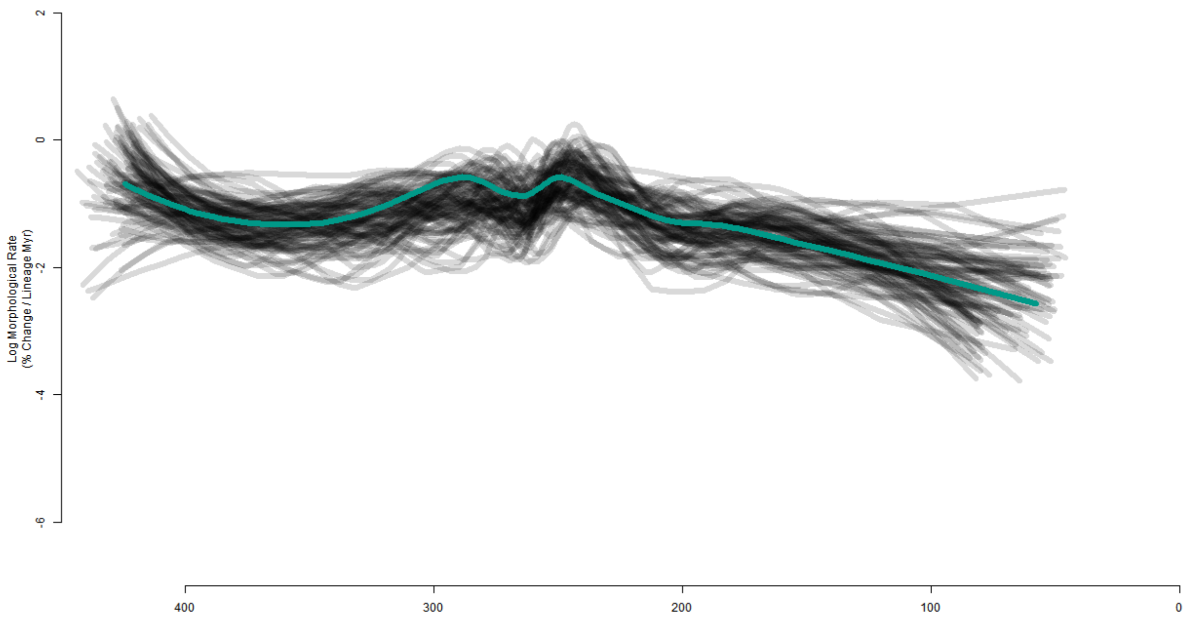

Log Morphological Rate
(% Change / Lineage Myr)
2
0
-2
-4
-6
400
300
200
100
0


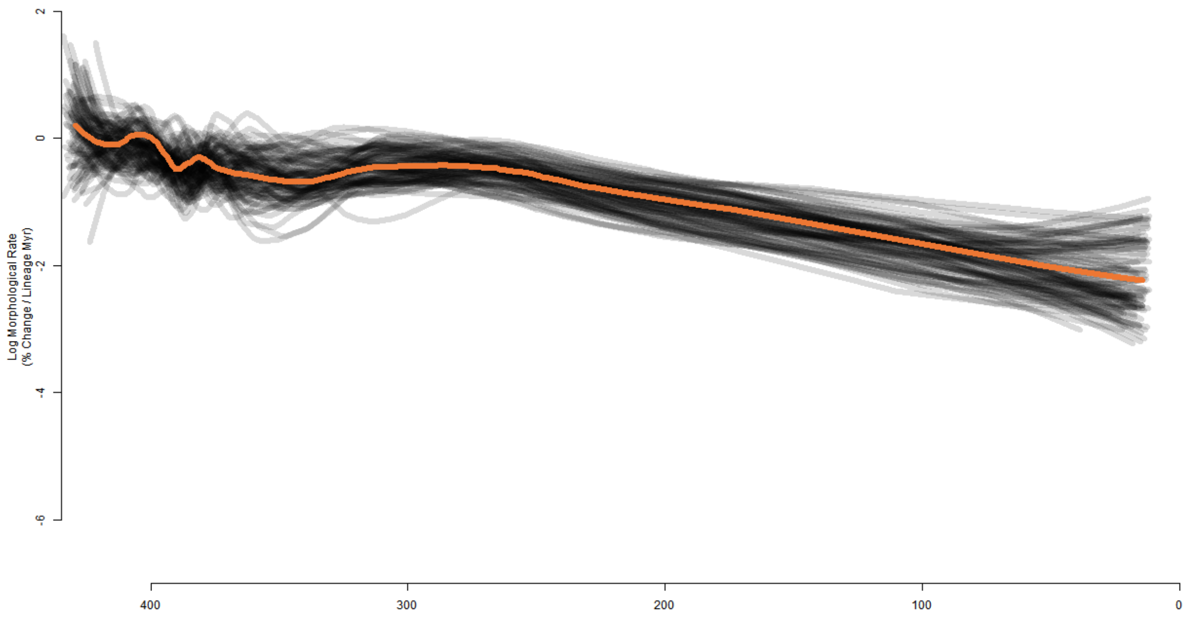

Log Morphological Rate
(% Change / Lineage Myr)
2
0
-2
-4
-6
400
300
200
100
0

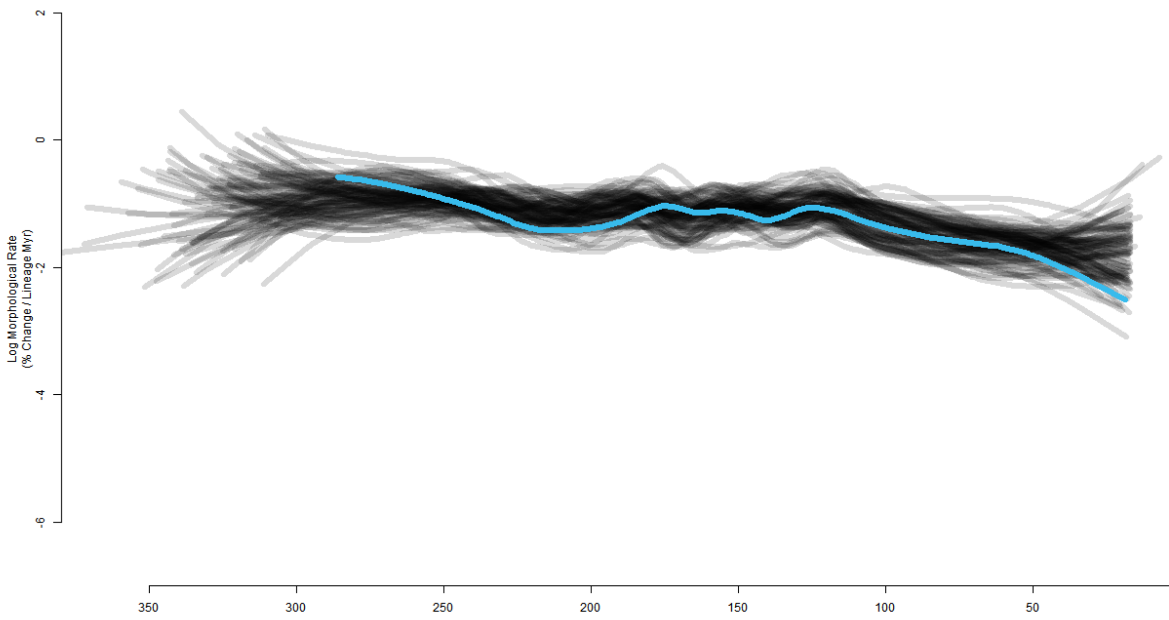


**Figure S9.** Average rate-through-time plots for discrete character evolution in coelacanths (top), lungfishes (centre), and holosteans (bottom). The solid coloured lines are the average rate across the MCC trees for each group. The semi-transparent grey lines behind the coloured lines are the average rates across a random sampling of 100 trees from the posterior. The averages are calculated and plotted at the midpoint of each branch in time, which results in the beginning and endpoints being offset from the root time and zero, respectively. All trees were generated using morphological data in BEAST v2.6.5 with the Fossilized Birth-Death model.

**Table S1.** Verbal summaries of results arising from analyses of three living fossil lineages.

| | **coelacanths** | **lungfishes** | **holosteans** |
|---|---|---|---|
| **discrete character rates through time** | strongly declining rates, with peak in the middle of clade history | strongly declining rates, with peak early in clade history | moderately declining rates, with relatively constant rates for much of clade history |
| **discrete character saturation** | minor saturation | strong saturation | moderate saturation |
| **body shape evolution** | overwhelming support for EB case of ACDC, with strongly declining rates over time | weak support for EB case of ACDC, with moderately declining rates over time | strong support for EB case of ACDC, with negligible decline in rates over time |

**Table S2.** *mvgls* model-fit GIC results for coelacanths using the MCC tree and the PC axes that summarised 100% of the variability. Models listed in the text. Fit = GIC scores, delta = change from highest-ranked score, w = GIC weights.

| | fit | delta | w |
|---|---|---|---|
| ACDC | -3,087.114 | 0 | 1 |
| BM | -3,013.844 | 73.270 | 0 |
| OU | -3,038.983 | 48.132 | 0 |

**Table S3.** *mvgls* model-fit GIC results for lungfishes using the MCC tree and the PC axes that summarised 100% of the variability. Models listed in the text. Fit = GIC scores, delta = change from highest-ranked score, w = GIC weights.

| | fit | delta | w |
|---|---|---|---|
| ACDC | -931.112 | 0 | 0.521 |
| BM | -928.540 | 2.572 | 0.144 |
| OU | -930.226 | 0.886 | 0.335 |

**Table S4.** *mvgls* model-fit GIC results for holosteans using the MCC tree and the PC axes that summarised 100% of the variability. Models listed in the text. Fit = GIC scores, delta = change from highest-ranked score, w = GIC weights.

| | fit | delta | w |
|---|---|---|---|
| ACDC | -28, 267.370 | 0 | 0.900 |
| BM | -28, 262.350 | 5.023 | 0.073 |
| OU | -28, 260.350 | 7.017 | 0.027 |

### References


1. Toriño P, Soto M, Perea D. 2021 A comprehensive phylogenetic analysis of coelacanth fishes (Sarcopterygii, Actinistia) with comments on the composition of the Mawsoniidae and Latimeriidae: evaluating old and new methodological challenges and constraints. *Hist. Biol.* **33**, 3423–3443. (doi:10.1080/08912963.2020.1867982)
2. Forey PL. 1998 *History of the Coelacanth Fishes*. Chapman & Hall.
3. Cavin L, Mennecart B, Obrist C, Costeur L, Furrer H. 2017 Heterochronic evolution explains novel body shape in a Triassic coelacanth from Switzerland. *Sci. Rep.* **7**, 13695. (doi:10.1038/s41598-017-13796-0)
4. Lloyd GT, Wang SC, Brusatte SL. 2011 Identifying heterogeneity in rates of morphological evolution: discrete character change in the evolution of lungfish (Sarcopterygii; Dipnoi). *Evolution* **66**, 330–348. (doi:10.1111/j.1558-5646.2011.01460.x)
5. Challands TJ, Smithson TR, Clack JA, Bennett CE, Marshall JEA, Wallace-Johnson SM, Hill H. 2019 A lungfish survivor of the end-Devonian extinction and an Early Carboniferous dipnoan radiation. *J. Syst. Palaeontol.* **17**, 1825–1846. (doi:10.1080/14772019.2019.1572234)
6. López-Arbarello A, Sferco E. 2018 Neopterygian phylogeny: the merger assay. *R. Soc. Open Sci.* **5**, 172337. (doi:10.1098/rsos.172337)
7. Latimer AE, Giles S. 2018 A giant dapediid from the Late Triassic of Switzerland and insights into neopterygian phylogeny. *R. Soc. Open Sci.* **5**, 180497. (doi:10.1098/rsos.180497)
8. Thies D, Waschkewitz J. 2016 Redescription of *Dapedium pholidotum* (Agassiz, 1832) (Actinopterygii, Neopterygii) from the Lower Jurassic Posidonia Shale, with comments on the phylogenetic position of *Dapedium* Leach, 1822. *J. Syst. Palaeontol.* **14**, 339–364. (doi:10.1080/14772019.2015.1043361)
9. Gibson SZ. 2016 Redescription and Phylogenetic Placement of †*Hemicalypterus weiri* Schaeffer, 1967 (Actinopterygii, Neopterygii) from the Triassic Chinle Formation, Southwestern United States: New Insights into Morphology, Ecological Niche, and Phylogeny. *PLOS ONE* **11**, e0163657. (doi:10.1371/journal.pone.0163657)
10. Xu G-H, Zhao L-J, Coates MI. 2014 The oldest ionoscopiform from China sheds new light on the early evolution of halecomorph fishes. *Biol. Lett.* **10**, 20140204. (doi:10.1098/rsbl.2014.0204)
11. Friedman M, Coates MI. 2006 A newly recognized fossil coelacanth highlights the early morphological diversification of the clade. *Proc. R. Soc. B Biol. Sci.* **273**, 245–250. (doi:10.1098/rspb.2005.3316)
12. Clarke JT, Friedman M. 2018 Body-shape diversity in Triassic–Early Cretaceous neopterygian fishes: sustained holostean disparity and predominantly gradual increases in teleost phenotypic variety. *Paleobiology* **44**, 402–433. (doi:10.1017/pab.2018.8)
